# A MOPD II-associated Pericentrin variant disrupts PACT domain dimerization and pericentriolar material recruitment

**DOI:** 10.64898/2026.05.01.722250

**Authors:** Makenzie S. Thomas, Brian J. Galletta, John M. Ryniawec, Anastasia Amoiroglou, Chaitali Khan, Carey J. Fagerstrom, Gregory C. Rogers, Nasser M. Rusan

**Author notes:** co-first authors (Makenzie S. Thomas, Brian J. Galletta).

## Abstract

Centrosome dysfunction is linked to developmental disorders affecting brain and body size, including microcephaly and primordial dwarfism. However, the cellular mechanisms underlying these rare conditions remain poorly understood. In this study, we investigate a rare variant of the centrosome-associated protein Pericentrin, which was discovered in a single family with Majewski/microcephalic osteodysplastic primordial dwarfism type II (MOPD II). Unlike the majority of pathogenic *PCNT* variants that cause severe protein truncation, the p.Lys3154del variant (ΔK3154) involves a single amino acid deletion in the protein’s only conserved functional domain, providing a unique opportunity to explore PCNT function in MOPD II. To model *PCNT*^ΔK3154^, we examined the effects of *Drosophila* Pericentrin-like protein (PLP) carrying an orthologous deletion (*Plp^ΔR^*). Our results show that *plp^ΔR^* animals exhibit smaller tissues that recapitulate MOPD II phenotypes. Behavioral assays revealed defects in climbing and mechanosensation, suggesting impaired sensory cilia function. We also found that *Plp^ΔR^* cells exhibit accelerated mitosis, increased apoptosis, and reduced pericentriolar material recruitment. *In silico* structural modeling, yeast two-hybrid, and co-immunoprecipitation experiments show that *Plp^ΔR^* produces a protein that disrupts PLP dimerization and PLP interaction with Asterless, another centrosome protein. Overall, modeling the human MOPD II patient variant *PCNT*^ΔK3154^ in *Drosophila* reveals how a single amino acid deletion affects biological processes from the molecular level to the organismal level. Our work offers new insights into the defective cellular mechanisms underlying MOPD II in patients with the *PCNT*^ΔK3154^ variant, potentially linking the etiology of the disease in these individuals to the loss of a single protein-protein interaction.

## Introduction

The centrosome is a dynamic, membraneless organelle that functions as the primary microtubule-organizing center (MTOC) in many eukaryotic cells (Gould and Borisy, 1977). Comprised of a pair of centrioles surrounded by a proteinaceous matrix called the pericentriolar material (PCM), the centrosome orchestrates several key cellular functions, including organizing the bipolar mitotic spindle, cilia formation, cell migration, and intracellular trafficking (Azimzadeh and Bornens, 2007; Bettencourt-Dias et al., 2011; Woodruff et al., 2014). PCM organization and accumulation at the centriole is often tightly coordinated with the cell cycle, peaking during mitosis to ensure robust microtubule nucleation, followed by a regulated disassembly upon mitotic exit (Nigg and Raff, 2009; Palazzo et al., 2000). The assembly, integrity, and function of the PCM relies on evolutionarily conserved scaffold proteins (Luders, 2012; Mennella et al., 2012; Varadarajan and Rusan, 2018), one of which is Pericentrin (PCNT).

PCNT is a large coiled-coil protein partially responsible for organizing PCM at the centrosome during mitosis (Dictenberg et al., 1998; Doxsey, 2005; Matsuo et al., 2010; Purohit et al., 1999; Takahashi et al., 2002). Germane to the function of PCNT is the ability to localize to the centrosome, at least in part through its highly conserved C-terminal pericentrin-AKAP450-centrosome targeting (PACT; Alderton et al., 2006) domain (Gillingham and Munro, 2000; Martinez-Campos et al., 2004). The PACT domain localizes to the centriole wall, while the N-terminus of PCNT extends radially outward into the PCM to anchor essential components of the centrosome such as γ-tubulin ring complexes, dynein, Chk1, and protein kinase A (PKA) (Dictenberg et al., 1998; Diviani et al., 2000; Lawo et al., 2012; Mennella et al., 2012; Purohit et al., 1999; Sorino et al., 2013). Mutations in *PCNT* that disrupt these processes have been linked to several human diseases, including cancer, ciliopathies, and several developmental disorders such as the autosomal recessive condition, Majewski/microcephalic osteodysplastic primordial dwarfism type II (MOPD II; MIM 210720) (Jurczyk et al., 2004; Majewski and Goecke, 1982; Majewski et al., 1982). Though MOPD II is the most common of the microcephalic primordial dwarfism syndromes, it is still quite rare, with only 150 cases identified worldwide as of 2017 (Bober and Jackson, 2017; Duker et al., 1993). The disorder is characterized by severe intrauterine and postnatal growth restriction, extreme short stature, microcephaly, skeletal dysplasia, distinctive craniofacial features, and cerebrovascular abnormalities (Fig. 1A) (Bober and Jackson, 2017; Bober et al., 2012; Duker et al., 1993; Majewski and Goecke, 1982; Majewski et al., 1982). Despite near-normal cognitive function, individuals with MOPD II can face complications such as insulin resistance, cardiac malformations, scoliosis, chronic kidney disease, and global vascular disease (Duker et al., 2021; Hall et al., 2004; Majewski and Goecke, 1982; Majewski et al., 1982; Rauch et al., 2008). Due to these associated comorbidities, patients with MOPD II often have a shortened life expectancy ranging from 7 to 41 years (Duker et al., 1993).

**Figure 1:**
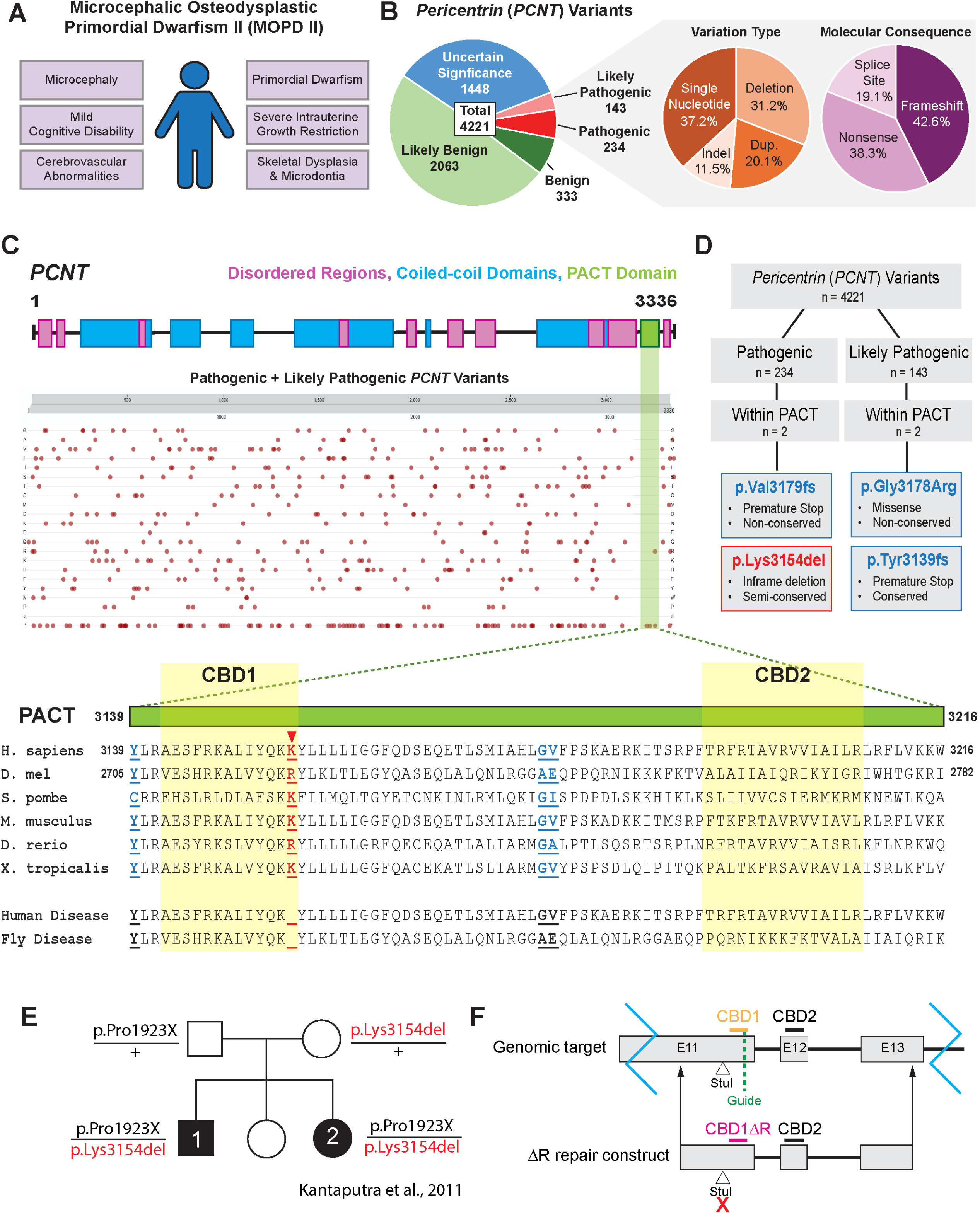
*PCNT* p.Lys3514del is a rare variant linked to MOPD II. **A.** Common clinical phenotypes of MOPD II. **B.** All reported *PCNT* variants categorized according to effect on human health. Pathogenic/Likely Pathogenic variants additionally categorized by variation type and molecular consequence. **C.** *PCNT* schematic highlighting the relative locations of disordered regions (pink), the PACT domain (green), and coiled-coil domains (blue). Pathogenic or likely pathogenic variants in *PCNT* are denoted in red dots by relative position (x-axis) and molecular change (y-axis; Rows 1-20: missense mutations by amino acid; Row 21: large deletions (d); Row 22: single amino acid deletions/frameshifts/premature stop codons (*)). Image is a screenshot from UniProt website (Gene ID: O96513 PCNT_HUMAN, “Variant viewer” tab. Enlarged PACT domain schematic scaled to align with evolutionary comparison of amino acid sequence with Human *PCN ^ΔK3154^* and Drosophila *Plp^Δ2720^* disease variant sequences denoted. Red arrowhead: conserved residue of interest, Lys3154. Blue: non-conserved variant residues. **D.** Flow chart of *PCNT* variant selection based on pathogenicity, location of variant within PACT domain, molecular consequence, and conservation. **E.** Pedigree of the family carrying the human PCNT variant, *PCNT^ΔK3154^* (Kantaputra et al., 2011; Kantaputra et al., 2004). **F.** Schematic of the generation of *Plp^ΔR2720^* using CRISPR. The top line represents the fly genome with E11, E12 and E13 being the last exons of the RF isoform of *Plp*. Calmodulin binding sites – CBD1, CBD2. The location of the guide RNA for cutting is indicated. The bottom line is the *Plp^ΔR2720^* repair construct that removes R2720 and introduces a silent change within a StuI site that was used for PCR-based screening.

The majority of reported MOPD II cases are caused by biallelic loss-of-function variants of *PCNT* (Duker et al., 1993). At the cellular level, MOPD II patient-derived cells or *PCNT* siRNA-treated cultured human cells show evidence of spindle abnormalities, chromosome misalignment or missegregation, premature sister chromatid separation, mitotic failure, and aneuploidy (Griffith et al., 2008; Rauch et al., 2008). Thus, many of the clinical manifestations reported in MOPD II are believed to stem from impaired PCM assembly and spindle pole integrity due to the loss of PCNT function, possibly leading to cell cycle defects, spindle organization and orientation defects, and ultimately an overall reduced cell count during development (Delaval and Doxsey, 2010).

Due to the limited clinical evidence available to guide the molecular understanding and treatment of rare diseases, modeling conditions such as MOPD II in tractable animal systems remains essential for uncovering the molecular basis of complex phenotypes and for the potential of identifying other effects of the variant allele that may have been missed as a result of small sample sizes seen in the clinic. *Drosophila melanogaster* provides a powerful platform for such studies. Approximately 60% of human genes have a counterpart in *Drosophila*, and about 75% of human disease-associated genes have a functional ortholog in flies (Fortini et al., 2000; Rubin et al., 2000; Ugur et al., 2016). Moreover, centrosome-associated diseases have broad functions across many cell types and give rise to diverse phenotypes, which can make them challenging to study in cell culture models. In contrast, *Drosophila* allows for the study of disease variants in the context of a whole animal, enabling cellular-level studies in multiple tissue types and developmental stages.

Pericentrin-like-protein (Plp), the *Drosophila* ortholog of PCNT, shares conserved functional domains and centrosome functions with human PCNT, including its roles in PCM assembly, mitotic spindle organization, and ciliogenesis (Martinez-Campos et al., 2004; Mennella et al., 2012). *Drosophila* Plp disruption leads to centrosome and ciliary phenotypes reminiscent of human disease, making it a compelling *in vivo* model for investigating pathogenic *PCNT* variants (Kawaguchi and Zheng, 2004; Martinez-Campos et al., 2004).

In this study, we model a rare MOPD II variant in *PCNT* using *Drosophila Plp* to investigate its impacts on the whole organism, individual tissues, cellular and subcellular functions, and protein interactions. We identified one particular variant, p.Lys3154del (ClinVar: ENST00000359568), as the only conserved mutation located within the PACT domain of PCNT that consists of a single, in-frame amino acid deletion (Kantaputra et al., 2004). By leveraging the behavioral, genetic, and imaging tools available in *Drosophila*, we aim to reveal mechanistic insights into how a rare pathogenic variant of *PCNT* compromises its function at the centrosome and contributes to MOPD II pathogenesis. We found that p.Lys3154del disrupts processes at the molecular, cellular, tissue, and organismal levels. Our work offers new insights into the failed cellular mechanisms underlying MOPD II, potentially linking this human disease to the loss of specific protein-protein interactions.

## Results

### Bioinformatic analysis of *PCNT* variants

To understand the current landscape of pathogenic *PCNT* variants, we consulted the OMIM, UniProt (2025), and ClinVar (Version: 2025-06-30) protein databases and found 4,221 *PCNT* variants (Henrie et al., 2018; Landrum et al., 2014). These variants were annotated based on their effects on human health (Fig. 1B). Of the 4,221 variants, 3,844 were classified as uncertain significance, likely benign, or benign (Fig. 1B). The remaining 377 variants are divided into pathogenic (234 variants) and likely pathogenic (143 variants). Of these pathogenic/likely pathogenic variants, 11.5% are indels, 20.1% are duplications, 31.2% are deletions, and 37.2% are single nucleotide substitutions (Fig. 1B). Molecularly, these 377 pathogenic/likely pathogenic variants include 19.1% splice site alterations, 38.3% nonsense mutations leading to premature protein truncation, and 42.6% frameshift mutations, with a significant number of these frameshifts resulting in premature protein truncation (Fig. 1B).

Pericentrin is a relatively unstructured protein containing many predicted coiled-coils and disordered regions (Fig. 1C). The majority of PCNT has limited amino acid sequence homology across species except for two small regions, one in the central region of the protein (HsPCNT: AA 1463-1502; DmPLP: AA 1523-1562), and the other is the C-terminal PACT domain. We were interested in investigating pathogenic/likely pathogenic variants that met three criteria: 1) occurs within the PACT domain, 2) does not result in protein truncation, 3) and the variant residue(s) are conserved in *Drosophila melanogaster*. Of four pathogenic/likely pathogenic variants within the PACT domain, only one met the criteria (Fig. 1D). The p.Lys3154del variant deletes three nucleotides resulting in the in-frame deletion of a single amino acid (ClinVar: ENST00000359568). This allele was identified in a single family of patients diagnosed with MOPD II (Fig. 1C-E) (Kantaputra et al., 2011; Kantaputra et al., 2004). The lysine residue deleted in this allele (HsPCNT: AA 3154) is either as a lysine (identical) residue or arginine residue (similar) across species from *S. pombe* to *D. melanogaster* (Fig. 1C). This suggests it could be essential for Pericentrin function, and therefore motivated our modeling of this variant in *Drosophila*.

Pericentrin-like-protein (PLP) is the *Drosophila* ortholog of human PCNT. While the overall sequence homology of PLP to PCNT is low (BLAST Global Alignment - PLP, PF isoform, vs. PCNT, NP_006022.3, 18% identity / 34% identity and similarity), they share several structural similarities, including multiple disordered regions, coiled-coil domains, the small conserved motif in the central region and the highly conserved PACT domain, which contains two Calmodulin-binding domains (CBD1 and CBD2; Fig. 1C) (Galletta et al., 2014; Kawaguchi and Zheng, 2004). In *Drosophila*, the analogous residue to PCNT Lys3154 is PLP Arg2720. Thus, to model the PCNT Lys3154 deletion (PCNT^ΔK^), we used CRISPR/Cas9 to edit the *Plp* locus to generate an Arg2720 deletion in *Drosophila Plp* that we refer to as *Plp^ΔR^* (Fig. 1F).

### Whole-animal characterization of *plp^ΔR^* mutants

We aimed to explore the effects of *Plp^ΔR^* in *Drosophila* as it related to MOPD II patient phenotypes, including microcephaly, dwarfism, and sensory deficits. As our interest was in understanding the nature of this variant allele, the majority of our experiments were performed in homozygous *Plp^ΔR/ΔR^* mutant flies. However, to control for the possibility of second site mutations on the *Plp^ΔR^* chromosome and to assess if this allele is a hypomorph by the classic genetic definition, most experiments were also performed with one *Plp^ΔR^* chromosome and a chromosome carrying a deletion that completely removes PLP and several surrounding genes (*Df(3L)Brd15*; thus the genotype *Plp^ΔR/Df^*). Furthermore, in many cases we have used a ubiquitously expressed PLP cDNA fused to GFP (*ubi-Plp::GFP*; (Galletta et al., 2014)) and a transgene containing a duplication of the genomic region surrounding *Plp* (*Plp^BAC^*) to demonstrate rescue of the phenotype. We hypothesized that homozygous *Plp^ΔR/ΔR^* mutant flies would present with phenotypes consistent with aspects of the presentation of MOPD II in patients and with previous studies investigating *PCNT* function (Miyoshi et al., 2009; Srsen et al., 2006; Wang et al., 2013; Zimmerman et al., 2004).

To begin, we performed a series of whole-animal phenotyping experiments examining changes in organ size, sensory function and fertility that one might expect to be associated with MOPD II and PCNT dysfunction. We found that *Plp^ΔR/ΔR^* and *Plp^ΔR/Df^* adult flies had smaller heads (Fig. 2A,B, S1A,B), both of which were rescued by the introduction of *ubi-Plp::GFP* or *Plp^BAC^* (Fig. 2A,B, S1C,D). We also found that *Plp^ΔR/ΔR^*, *Plp^ΔR/Df^*, and null (*Plp^2172/Df^*) mutant flies all had significantly decreased adult wing area (Fig. 2C, D, Fig. S2A,B), which were also rescued by the introduction of *ubi-Plp::GFP* or *Plp^BAC^* (Fig. 2C,D, S2C,D). As there is an established strong linkage between mutations in genes encoding centriole and centrosome proteins and smaller tissues, we explore this wing size phenotypes in more detail.

**Figure 2:**
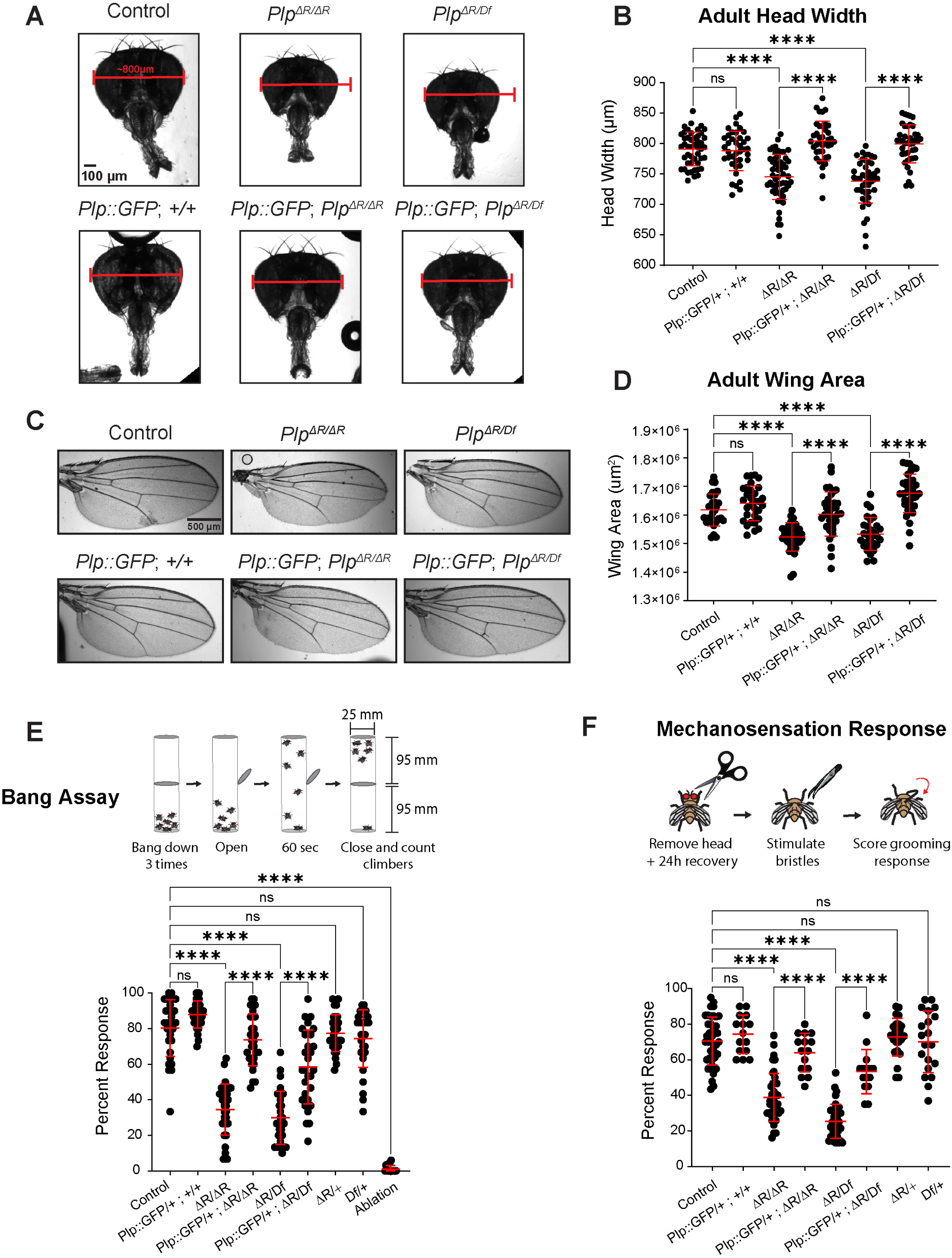
Adult tissue characterization of *Plp^ΔR^* mutants. **A.** Adult heads. 800 µm bars for ease of comparison. **B.** Adult head width measurements. Additional genotypes in Figure S1. **C.** Adult wings. **D.** Wing area (µm^2^) measurements taken by tracing the margin of the wing from vein L1 to L6. Additional genotypes in Figure S2. **E.** Schematic of bang assay. The percentage of flies that climbed to upper chamber within 60s is reported. Each point represents the average climbing response from a group of 15-30 flies assayed three times. Ablation = Control fly with arista surgically removed to impair graviception. **F.** Schematic of mechanosensory assay. Percentage of front leg grooming response is reported. Each point represents the average grooming response from a group of 10 flies assayed three times. Numbers of samples measured are in Supplementary File 1. n.s. = not significant, **** = p ≤ 0.0001.

Of note, we observed two phenotypes unique to the homozygous *Plp^ΔR/ΔR^* flies — missing and shortened bristles at the wing margin and a shorter distance between veins L3 and L4 (Figure S2A). However, these phenotypes were not rescued by exogenous PLP, indicating that they are a consequence of other unknown mutations on the *Plp^ΔR^* chromosome; we did not explore these phenotypes further.

In addition to roles in centrosomes, PCNT plays important roles in ciliogenesis. In human cell culture and in mice, PCNT localizes to the base of cilia (Jurczyk et al., 2004; Majewski and Goecke, 1982; Majewski et al., 1982), and in patient-derived fibroblasts and cultured cells where PCNT has been silenced, there is a loss of primary cilia (Jurczyk et al., 2004; Stiff et al., 2016). *PCNT* mutant mice have defects in the assembly of olfactory cilia, migration of interneurons in the olfactory bulb, and show defects in olfactory function (Chen et al., 2019; Endoh-Yamagami et al., 2010; Guirgis et al., 2001; Muhlhans et al., 2011). Interestingly, cilia in motile and non-neuronal epithelia in *PCNT* mutant mice appear unaffected (Chen et al., 2019; Endoh-Yamagami et al., 2010; Guirgis et al., 2001; Muhlhans et al., 2011). While MOPD II patients do not present many of the classic phenotypes associated with ciliopathies, some MOPD II patients have ocular and kidney issues, which raise the possibility of cilia involvement (Bober et al., 2010; Chen et al., 2019; Duker et al., 2021; Endoh-Yamagami et al., 2010; Guirgis et al., 2001; Muhlhans et al., 2011), although whether these issues arise specifically from cilia dysfunction has not been examined.

There is a clear link between PLP and cilia function in *Drosophila* as *Plp* null flies have severe defects in ciliated sensory organ structure and function (Galletta et al., 2014; Martinez-Campos et al., 2004). We therefore tested whether *Plp^ΔR^* flies present with sensory defects using two behavioral assays. We performed a “bang assay” to assess the ability of flies to climb upward after a strong downward force (Figure 2E). Flies unable to sense gravity, because of surgical removal of the arista from their antenna (Fig. 2E. ablation), do not climb in this assay. We observed a severe deficit in this assay in *Plp^ΔR/ΔR^* and *Plp^ΔR/Df^* flies, with a 57.1% and 62.8% decrease, respectively, compared to controls, which could be rescued by Plp::GFP (Figure 2E). While this assay can assess graviception, flies with locomotor or proprioceptive defects might also fail to climb. We therefore performed the same assay, but placed a light source above the apparatus (Figure S3A). Interestingly, *Plp^ΔR/ΔR^* flies also failed to climb in this assay (Figure S3B). While this confirms that the flies have a climbing deficit, the assay still requires the flies to move against gravity and flies that cannot sense gravity (ablation) do not perform as well as controls. The fact that we see a further reduction of climbing in the phototaxis assay when the arista is removed from *Plp^ΔR/ΔR^* flies.suggests that they may have an underlying gravitaxis defect that exacerbates their more general climbing deficit. To test mechanosensation more directly, we performed a classical assay of sensory bristle function in which the notopleural or postalar bristles of decapitated flies were mechanically stimulated and the percentage of flies that displayed grooming behavior was recorded (Figure 2F). We observed severe response deficits in both *Plp^ΔR/ΔR^* and *Plp^ΔR/Df^* with a 45.0% decrease and 64.1% decrease, respectively, compared to controls (Figure 2F), which could be rescued by *Plp::GFP*. Combined, these results demonstrate that *Plp^ΔR^* causes climbing and sensory defects, which could result from altered sensory organ structure, impaired sensory cilia formation, or errors in processing sensory information in the brain.

Centrosomes also contribute to the formation of viable sperm where they are required for meiosis and for linking the sperm head to the tail. While male infertility has not been reported in patients with MOPD II, prior studies have highlighted the importance of PLP in sperm development. For example, *Plp* null flies have defects in centriole orientation and positioning in pre-meiotic spermatocytes, fail to produce motile sperm and have seminal vesicles devoid of sperm (Galletta et al., 2014; Galletta et al., 2020; Martinez-Campos et al., 2004)(Roque et al., 2018). However, *Plp^ΔR/ΔR^* flies can be maintained as a healthy, robust stock suggesting that they do not have significant defects in mating or fertility.

### Examination of underlying tissues in *Plp^ΔR^* mutants

We next examined *Plp^ΔR^* flies for tissue-level phenotypes typically associated with centrosome and cilia dysfunction. Since we had seen an effect on head size, we examined the adult brain size in *Plp^ΔR^* flies using whole-brain volumetric analysis (Fig. 3A). We found that flies reared at 15°C or 25°C showed no difference in brain volume (Fig. 3B). This suggests that our assay for brain size is either 1) unable to identify brain size variation for a technical reason, for example because of a fixation artefact, or 2) the underlying cause of the small head phenotype might not be a result of a brain size difference. This lack of effect on brain size is in contrast to other microcephaly genes studied in *Drosophila* that have reported smaller brains (Poulton et al., 2017; Ramdas Nair et al., 2016; Robinson et al., 2020; Schoborg et al., 2015). The origin of this difference could reveal differences in the cellular consequence of mutations in these genes. Due to the lack of a clear brain size phenotype in *Plp^ΔR^* flies, we did not pursue analysis of brains at the cellular level.

**Figure 3:**
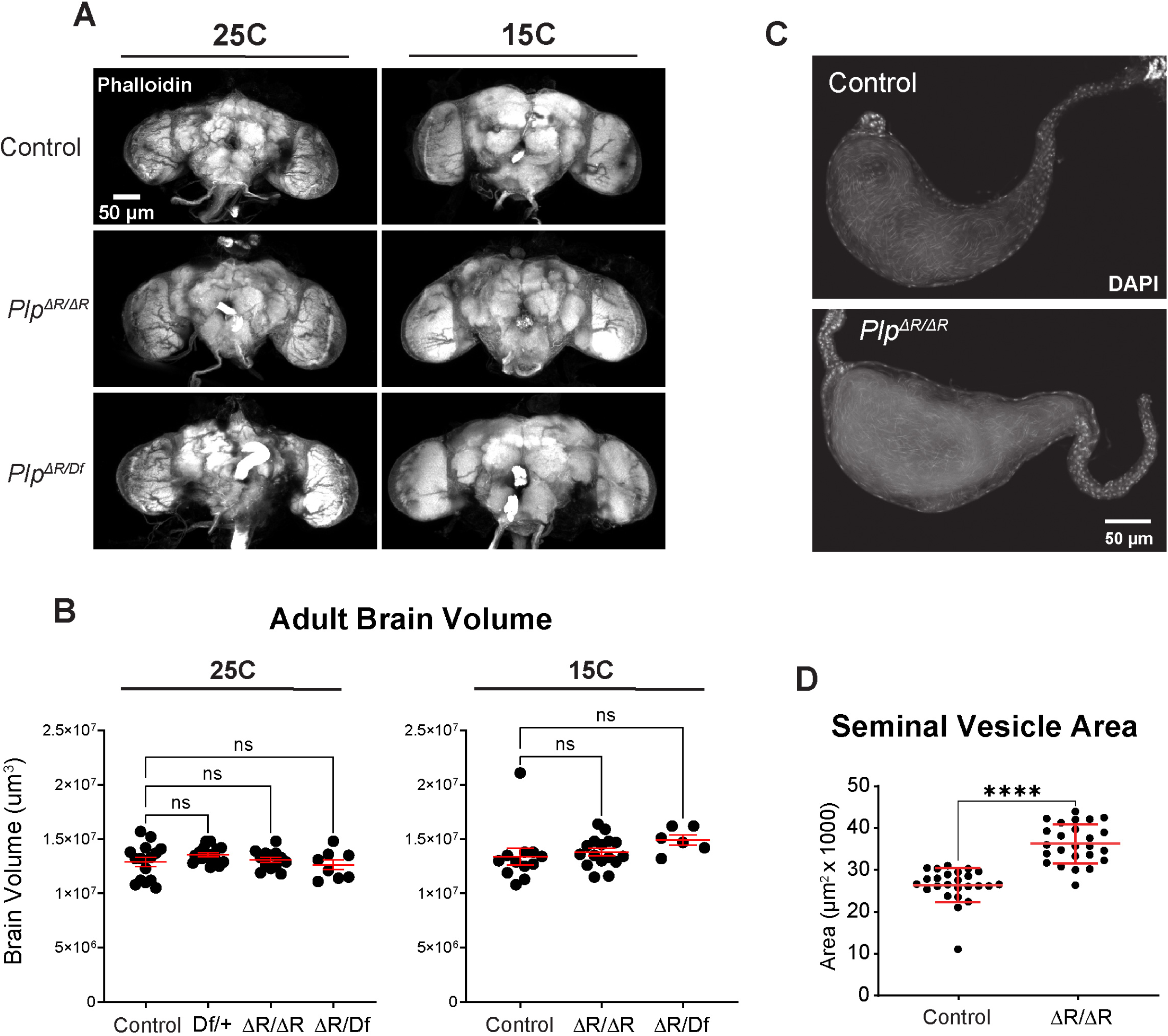
Tissue-level characterization of *Plp^ΔR^* mutants. **A.** 3D projections of adult fly brains stained with phalloidin reared at 25°C and 15°C. **B.** Brain volume (µm^3^) measurements from flies reared at 25°C and 15°C. **C.** Seminal vesicles from naïve males stained with DAPI. **D.** Seminal vesicle area (µm^2^) measurements. Numbers of samples measured are in Supplementary File 1. n.s. = not significant, **** = p ≤ 0.0001.

To determine if the fertility we observe in the *Plp^ΔR/ΔR^* stock might hide an underlying defect in sperm development, insufficient to affect organismal fertility, we examined seminal vesicles from adult males and found that *Plp^ΔR/ΔR^* seminal vesicles contained sperm (Fig. 3C). We also measured seminal vesicle cross-sectional area as a proxy for the production of functional sperm, and observed a significant increase in *Plp^ΔR/ΔR^* males compared to controls (Fig. 3D). These results indicate that *Plp^ΔR^* does not impair spermatogenesis, although the increased area could indicate an increase in the rate of sperm production. The qualitative fecundity of the homozygous *Plp^ΔR/^*^ΔR^, and the robust number of sperm in the seminal vesicles indicate that *Plp^ΔR^* does not significantly impair spermatogenesis or sperm function. Given the lack of a clear negative effect on fertility, we did not pursue further analyses of sperm development in this study.

To examine sensory neurons, we investigated the Johnston’s Organ (JO), which contains an array of ciliated sensory neurons that are used to sense gravity, among other functions (Fig. S4A, B). JO sensory neurons of *Plp* null flies lack normal dendritic extensions (Martinez-Campos et al., 2004) and have mispositioned centrioles are no longer located at the interface between the dendrite of neuron and Scolopale cells, where cilia are located (Galletta et al., 2014). Additionally, PLP protein that cannot bind calmodulin due to mutations in CBD2 does not localize to the centrioles of the JO (Galletta et al., 2014). We hypothesized that *Plp^ΔR^* might exhibit a similar defect in the formation or positioning of the mechanosensory cilia. However, we found that the JO centrioles of *Plp^ΔR/ΔR^* flies were qualitatively normal, showing properly positioned centrioles and normal PLP localization (Figure S4C). While the lack of a severe phenotype in the JO was qualitatively clear, we could not determine if there was a subtle effect of PLP levels within the JO because of technical limitations on quantitative analysis of immunofluorescence signals in this tissue. Thus, the cellular cause of mechanosensation and climbing defects remains to be explored.

### Plp^ΔR^ negatively impacts mitosis and increases cell death in developing tissues

The smaller wing size we observed in *Plp^ΔR/ΔR^* is consistent with the severe intrauterine growth restriction and dwarfism that are defining clinical phenotypes of MOPD II. It has been previously shown that cells derived from MOPD II patients or human cell lines depleted of PCNT have mitotic defects, aneuploidy, and increased cell death (Griffith et al., 2008; Rauch et al., 2008; Zimmerman et al., 2004). Thus, we hypothesized that *Plp^ΔR^* may impact cell division or promote apoptosis in the wing imaginal discs, the precursor larval tissue from which wings develop during pupation. The relatively flat morphology of these tissues allows for easy identification, immunostaining, and live imaging of mitotic and apoptotic cells.

To test the impact of *Plp^ΔR^* on proliferation, we measured wing disc mitotic index (phospho-histone H3 positive cells per wing disc pouch; Fig. 4A; S5A) and observed a significant decrease in the percentage of mitotic cells in *Plp^ΔR/ΔR^*, *Plp^ΔR/Df^*, and *Plp^2172/Df^*, with *Plp^2172/Df^* null wing discs displaying the strongest phenotype (Fig. 4B, S5B). Importantly, the reduction in mitotic index was rescued by *Plp::GFP* and *Plp^BAC^*. There are at least three reasons why the mitotic index might be reduced in *PLP^ΔR^*: 1) mutant wing disc cells have an extended interphase, thus, mitosis is a smaller percentage of the overall cell cycle, 2) the duration of mitosis is reduced, or 3) there are fewer cells with proliferative potential, possibly as a result of proliferating cells undergoing cell death. We were not able to examine interphase duration given its length in this tissue, but we did measure mitotic duration by live cell imaging. We found that the time from nuclear envelope breakdown (NEB) to anaphase-onset (AO) was slightly reduced in *Plp^ΔR/ΔR^* cells (Control = 11.3 ± 0.71 minutes; *Plp^ΔR/ΔR^* = 9.9 ± 0.74 minutes; Fig. 4C, D). This is consistent with the reduction in mitotic index, but does not eliminate other factors contributing to this reduction.

**Figure 4:**
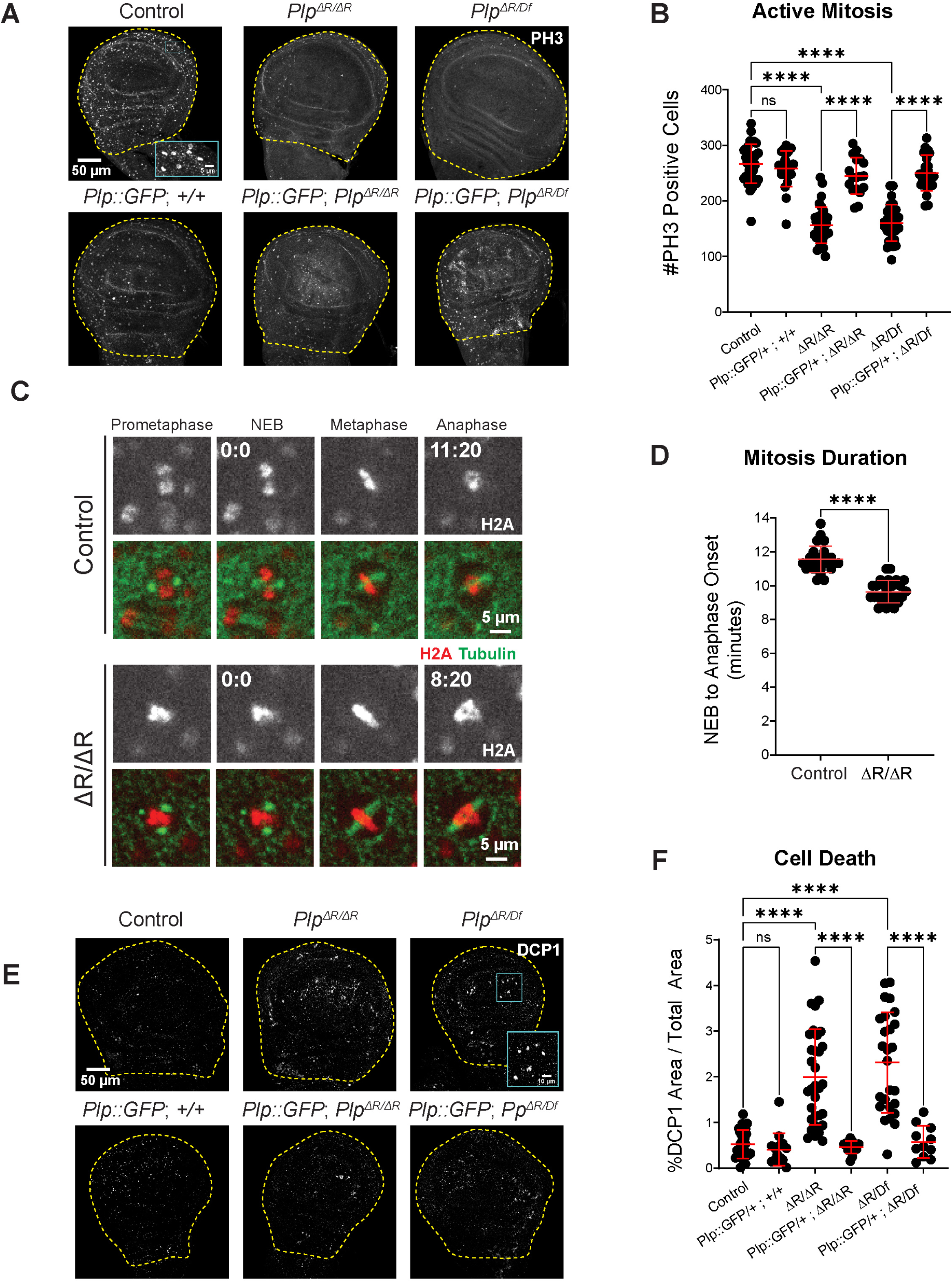
*Plp^ΔR^* alters mitosis and increases cell death. **A.** Wing imaginal discs from wandering third instar larvae stained for DNA (DAPI, blue) and active mitosis (phospho-histone H3 (PH3)). The combined wing pouch and hinge region are outlined. Inset: examples of PH3-positive cells. **B.** Number of PH3-positive nuclei in total wing pouch and hinge area. Additional genotypes in Figure S5A, B. **C.** GFP::tubulin and RFP::H2A in wing disc epithelial cells undergoing mitosis. **D.** Time (minutes) measured from nuclear envelope breakdown (NEB) to anaphase onset (AO) in wing disc epithelial cells. **E.** Wing imaginal discs from wandering third instar larvae stained for DNA (DAPI, blue) and apoptosis (cleaved *Drosophila* Death caspase-1 (DCP1)). The combined wing pouch and hinge region are outlined. Inset: examples of DCP1-positive cells. **F.** Percentage of DCP1-positive area out of total wing pouch and hinge area. Additional genotypes in Figure S5E, F. Numbers of samples measured are in Supplementary File 1. n.s.= not significant, ****p ≤ 0.0001.

Interestingly, even though these wing disc cells appear to rush through mitosis, we did not detect any gross spindle abnormalities or obvious chromosome segregation errors such as lagging chromosomes or DNA bridges. We also did not see an increase in double stranded DNA breaks (DSBs) in *Plp^ΔR/ΔR^ or Plp^ΔR/Df^* wing discs by staining for phosphorylated histone 2A variant (γH2Av), although we did observe a significant increase in DSBs in *Plp^2172/Df^*. (Figure S5C, D) Furthermore, we did not observe an immediate transition to cell death from mitosis as cells appeared to fully exit mitosis into interphase. It is worth noting that these cells are extremely small, making it difficult to detect small but significant mitotic errors such as Ultrafine Anaphase Bridges (UFBs), which can ultimately trigger cell death (Fernandez-Casanas and Chan, 2018; Liu et al., 2014).

To determine if *Plp^ΔR^* wing cells experience higher rates of cell death, we immunostained wing discs for cleaved Death caspase-1 (DCP1) and found that *Plp^ΔR/ΔR^*, *Plp^ΔR/Df^*, and *Plp^2172/Df^* all displayed increased apoptosis compared to controls (Fig. 4E,F; S5E,F), which can be rescued by *Plp::GFP*. Thus, a simple model is that *Plp^ΔR^* results in DNA damage that triggers cell death, possibly as a result of an accelerated mitosis that bypasses mitotic checkpoints (Basu et al., 1999; Sorino et al., 2013; Taylor and McKeon, 1997; Tibelius et al., 2009). This simple model would also explain the reduction in adult wing area (Fig. 2D). An alternative model is that the cell death and accelerated mitoses phenotypes in *Plp^ΔR^* are independent (see discussion).

After observing the increased cell death in wing discs, we were curious if cell death occurred in other imaginal discs. We therefore performed immunostaining for DCP1 in the leg and eye discs and found a similar increase in apoptosis for *Plp^ΔR/ΔR^*, *Plp^ΔR/Df^*, and *Plp^2172/Df^* compared to controls (Figure S6). This suggests that loss of R2720 in PLP may lead to global defects in epithelial tissue development, which is consistent with the overall small stature of MOPDII patients.

### *plp^ΔR^* impacts PCM recruitment

Given the deleterious impact of *Plp^ΔR^* on cell viability, tissue morphogenesis, and overall sensory defects, we next sought to use our disease model to understand the molecular changes occurring in wing disc cells. PLP is well characterized as a component of the centrosome, residing on the surface of the centriole in a region termed the “bridge zone” (Fu and Glover, 2012; Lawo et al., 2012; Mennella et al., 2012; Sonnen et al., 2012; Varadarajan and Rusan, 2018). We previously demonstrated that alterations in Calmodulin Binding Domain 2 (CBD2) within the PACT domain significantly reduces PLP localization to the centrosome (Galletta et al., 2014). We therefore hypothesized that *Plp^ΔR^*, which is a mutation in Calmodulin Binding Domain 1 (CBD1; Fig. 1C), would also disrupt PLP recruitment to the centrosome.

Using quantitative immunofluorescence in wing discs, we found *Plp^ΔR/ΔR^* had an approximately 35% reduction in PLP levels at metaphase centrosomes compared to control metaphase cells (Fig. 5A,C). Further protein level reduction was seen in hemizygous *Plp^ΔR/Df^* and null *Plp^2172/Df^*, which have an approximately 80% and 95% reduction in PLP levels, respectively. During interphase, *Plp^ΔR/ΔR^* and *Plp^ΔR/Df^* exhibited PLP protein level decreases at the centriole of approximately 45% and 60% respectively (Fig. S7A), indicating that the reduced levels of PLP^ΔR^ localization are not specific to mitosis. This eliminates the possibility that the reduction in PLP^ΔR^ localization is due to loss of PLP interaction with PCM proteins, which accumulate dramatically on centrosomes during mitosis in *Drosophila* (Rogers et al., 2008). Finally, we performed Western blots on wing discs from wandering 3^rd^ instar larvae and measured a 70% decrease in PLP levels in *Plp^ΔR/ΔR^* (Figure S7B-D). Thus, at this time we cannot determine if the reduction of *Plp^ΔR/ΔR^* at the centriole is a result of less total protein, less centriole binding, or both. Detailed biochemical and protein turnover studies will be required to untangle the contributions of these factors to the final amount of PLP on the centriole.

**Figure 5:**
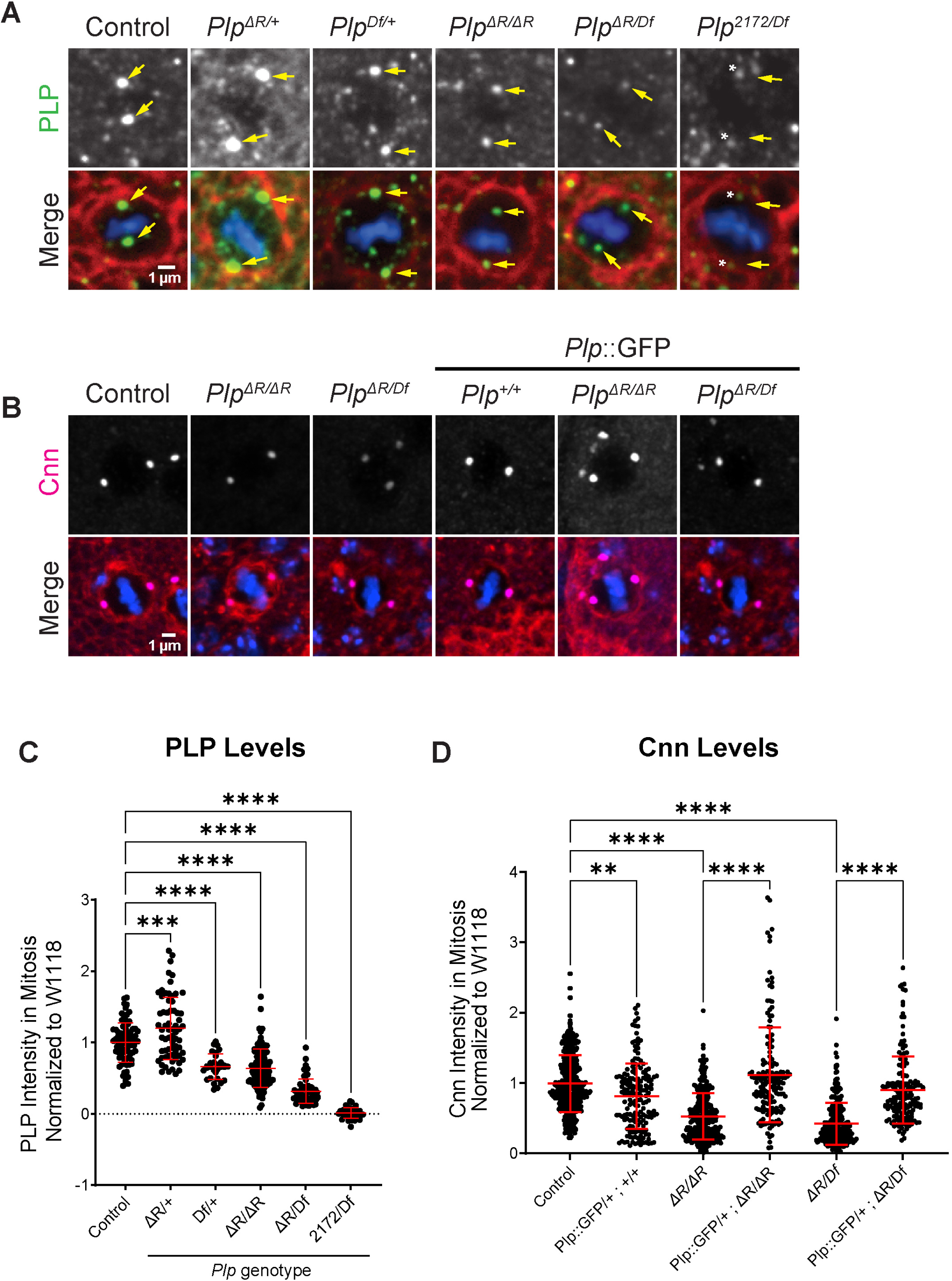
*Plp^ΔR^* impacts PCM recruitment. **A.** Metaphase wing disc epithelial cells stained for PLP (green), phalloidin (red) and DAPI (Blue). Yellow arrows: location of centrosomes. Asterisks indicate spots in the PLP channel that do not colocalize with centriole markers (not shown). **B.** Metaphase wing disc epithelial cells stained for PCM component, Centrosomin (Cnn, magenta), phalloidin (red) and DAPI (blue). **C.** PLP intensity at metaphase centrosomes normalized to *w^1118^* controls. **D.** Cnn intensity at metaphase centrosomes normalized to *w^1118^* controls. Numbers of samples measured are in Supplementary File 1. **p ≤ 0.01, ***p ≤ 0.001, ****p ≤ 0.0001.

Previous studies in neuroblasts, primordial germ cells, and precellularized embryos have shown that *Plp* null mutants have defects in PCM recruitment (Galletta et al., 2014; Lerit et al., 2015; Ma and Viveiros, 2014; Martinez-Campos et al., 2004). (Fang and Lerit, 2020). To test if PCM levels were affected in *Plp^ΔR^* centrosomes, we performed quantitative immunofluorescence of Centrosomin (Cnn) at metaphase centrosomes in wing discs. We found that *Plp^ΔR/ΔR^, Plp^ΔR/Df^*, and *Plp^2172/Df^* metaphase centrosomes have a significant reduction in Cnn, which is fully rescued by *PLP::GFP and Plp^BAC^* (Fig. 5B,D; Figure S8).

One critically important detail in our analysis is that loss of a single copy of *Plp* (*Plp^Df/+^*) reduces PLP levels at metaphase centrosomes (Figure 5C), but *does not* result in Cnn reduction (Fig. S8). In contrast, *Plp^ΔR/ΔR^*, which results in a reduction in PLP levels comparable to *Plp^Df/+^* (Fig. 5C), *does* result in a significant reduction in Cnn (Fig. 5D). This indicates that the decrease in Cnn at metaphase centrosomes is not solely a consequence of reduced PLP levels, but rather reflects a specific, direct effect of the R2720 deletion in PLP^ΔR^ on PCM levels.

### *Plp^ΔR^* disrupts secondary and tertiary protein structure

To better understand how the *Plp*^ΔR^ mutation might affect the function of the PLP protein, we utilized *in silico* methods to probe the possible structure of wild-type and mutant PLP. Much of the PLP protein is predicted to consist of regions that are unstructured and others that could form coiled-coils, including within the PACT domain (Fig. 1C). Using WaggaWagga, a coiled-coil prediction server (Simm et al., 2015; Simm et al., 2021), we investigated if the PLP^ΔR^ mutation affects the probability of coiled-coil formation within PACT (DmPACT; AA 2634-2780). We found that there is a significant likelihood that the N-terminal α-helix in DmPACT could form a coiled-coil up to CBD2 (Fig. 6A; AA 2645-2746). Similar analysis revealed that deletion of R2720 alters the register of the α-helix, reducing the predicted coiled-coil to only residues 2641-2684, several residues before CBD1 (Fig. 6A’).

**Figure 6:**
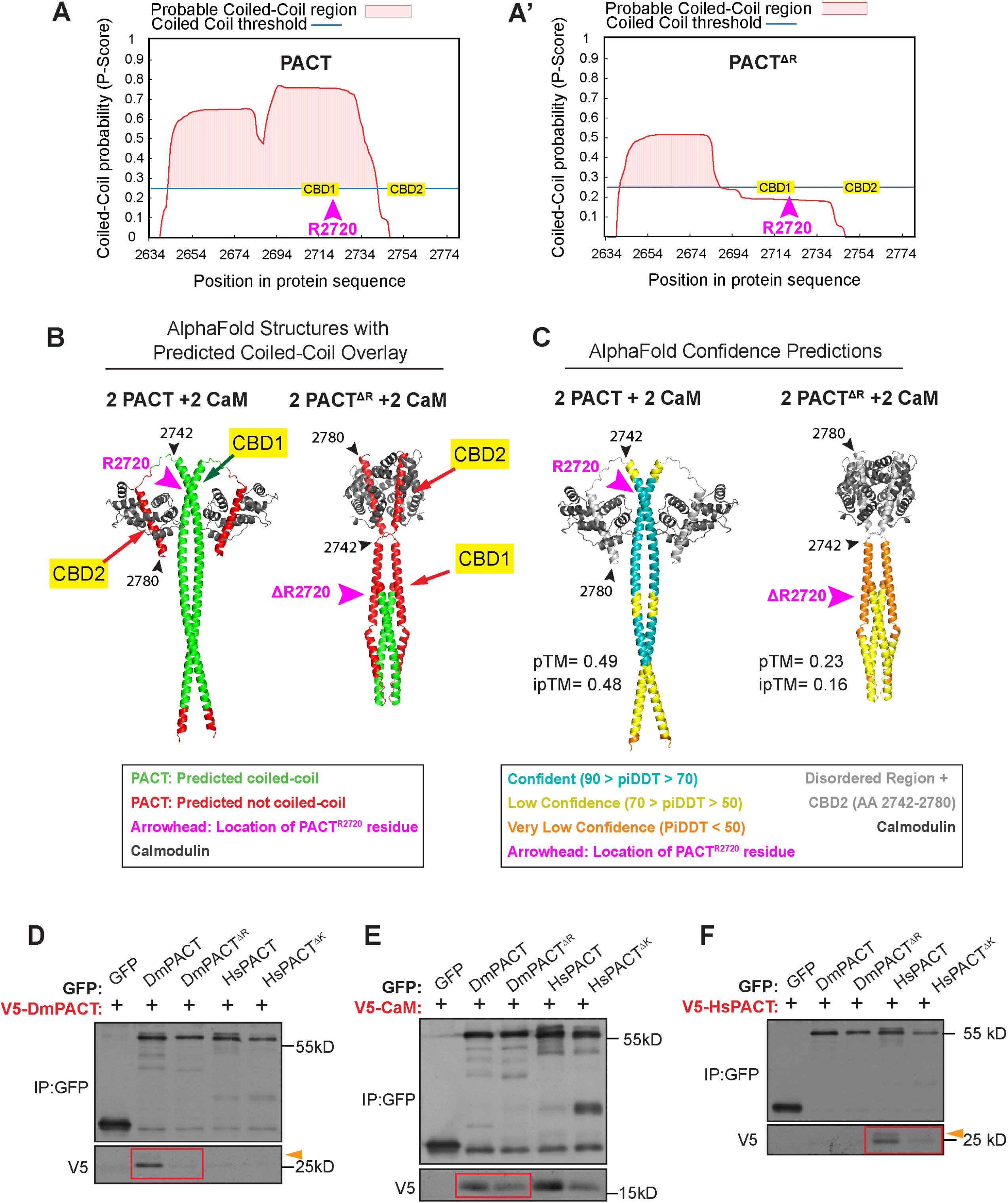
*Plp^ΔR^* disrupts PACT dimerization. **A.** Coiled-coil predictions of DmPACT domain from Waggawagga. CBD1 and 2 are indicated as is the position of R2720. **A’.** Coiled-coil predication of DmPACT^ΔR^ domain. **B.** AlphaFold3 structural predictions of 2 DmPACT^WT^ + 2 Calmodulin and 2 DmPACT ^ΔR^ + 2 Calmodulin overlayed with colors to represent predicted coiled-coil regions (green) and regions not predicted to form coiled-coils (red). R2720 in WT and its expected position in the ΔR are indicated by arrows (pink). Residues 2742 and 2780 (black arrows) are used as landmarks to highlight the mispositioning of CBD2 in DmPACT^ΔR^. **C.** Same structures as B. Color overlay corresponds to AlphaFold3 confidence scores. R2720 in WT and its expected position in the ΔR are indicated by arrows (pink). Residues 2742 and 2780 (black arrows) are used as landmarks to highlight the mispositioning of CBD2 in DmPACT^ΔR^. **D.** Co-immunoprecipitation is S2 cells showing V5-DmPACT interaction with GFP-DmPACT (red box, left), but not with DmPACTΔR (red box right), HsPACTWT, or HsPACTΔK. Arrowhead highlights slower migrating species. **E.** Co-immunoprecipitation is S2 cells showing V5- CaM interaction with both DmPACTWT and DmPACTΔR (red box). V5- CaM also interacts with HsPACTWT, and HsPACTΔK **F.** Co-immunoprecipitation is RPE cells showing V5-HsPACT interaction with GFP-HsPACT, but not with HsPACTΔK, DmPACTWT, or DmPACTΔR. Arrowhead highlights slower migrating species.

We next utilized AlphaFold3 to visualize how the PLP^ΔR^ mutation might affect protein folding. We modeled DmPACT^WT^ and DmPACT^ΔR^ as monomers, heterodimers, homodimers, and oligomers with and without Calmodulin (Fig. S9). The DmPACT^WT^ homodimer is predicted, with high-confidence, to form a parallel coiled-coil structure that is separated from the C-terminal CBD2 domains by an unstructured region, consistent with the WaggaWagga results (Fig. 6B, S9C). This structure suggests flexibility between the main coiled-coil and CBD2 which would allow for Calmodulin (CaM) binding (Fig. S9C). The predicted DmPACT^WT/ΔR^ heterodimer indicates significant disruption in the ability to form a coiled coil with much lower overall confidence, while the predicted DmPACT^ΔR^ homodimer resulted in an even lower confidence structure (Fig. S9C). These results are consistent with the WaggaWagga results, as AlphaFold predicts a short coiled-coil region near the N-terminus of the DmPACT^ΔR^ homodimer that is significantly shorter than the DmPACT^WT^ homodimer (Fig. 6A’, C). These predictions suggest that deletion of R2720 disrupts the long α-helix in PACT, altering the molecular distance between amino acids and, in turn, preventing formation of the putative coiled-coil structure of the PACT dimer.

To test the *in silico* models of PACT dimerization in cells, we next performed co-immunoprecipitations (co-IPs) of fly and human PACT constructs transiently expressed in cultured *Drosophila* S2 cells. Anti-GFP IPs were performed from cells co-transfected with GFP- and V5-tagged PACT expression constructs. We found that GFP-DmPACT robustly co-IPed V5-DmPACT; however, despite their similar predicted structures, the human PACT ortholog (HsPACT) did not co-IP with DmPACT (Fig. 6D; S10A). As predicted by Alphafold3, deletion of R2720 disrupted the association with wild-type DmPACT (Fig. 6D, S10A).

Lastly, we modeled the homo and heterodimers of PACT with two CaM proteins. The predicted structure shows that the CBD2 domains would facilitate CaM binding with high-confidence scores. In contrast, CBD1, which is modeled as buried within the coiled-coil, likely does not bind CaM (Fig. 6B, C). Modeling four CaM proteins with dimerized PACT corroborated this prediction (not shown). When modeling the DmPACT^ΔR^ homodimer with two CaMs, the Alphafold3 model predicts a very low confidence, grossly misfolded structure, with the N-terminus of PACT losing most of its coiled-coil structure and folding on itself (Fig. 6B,C). However, the predicted DmPACT^ΔR^ homodimer model is still able to bind CaM via CBD2, which we confirmed by co-IP experiments in cultured S2 cells using GFP-tagged PACT^ΔR^, from both flies and humans, and V5-tagged CaM; although PACT^ΔR^ shows reduced CaM binding compared to wild-type PACT (Fig. 6E; S10B, C). This interaction is consistent with our previous study of PLP-CaM where we showed that CBD2 is the main CaM binding domain (Galletta et al., 2014). Overall, deletion of R2720, which resides in CBD1, blocks PACT dimerization and reduces CaM binding. Our data suggests that CBD1 does not physically bind CaM, but instead, indirectly influences the CBD2-CaM interaction through CDB1’s role in PACT dimerization. Possibly, loss of dimerization results in the inability of PLP^ΔR^ to efficiently bind other centrosome partners for efficient localization or PCM recruitment.

To investigate our *Drosophila* protein findings in human PCNT, we performed similar AlphaFold3 analysis of the human PACT (HsPACT^WT^) and HsPACT^ΔK^, which revealed nearly identical predictions to DmPACT (Fig. S11) where coiled-coil formation and dimerization are disrupted in HsPACT^ΔK^ and CaM binding is restricted to CDB2. Furthermore, we confirmed HsPACT dimerization as GFP-HsPACT co-IPed V5-HsPACT, and showed that deletion of K3154 disrupted dimerization (Fig. 6F; S10D). Finally, we found that, similar to *Drosophila*, CaM binding is reduced in human HsPACT^ΔK^ (Fig. 6E).

### PLP^ΔR^ loses protein-protein interaction with Asterless

PCNT is a large coiled-coil protein predicted to form oligomeric structures, including dimers and other higher-order assemblies necessary to recruit and organize PCM (Dictenberg et al., 1998; Gillingham and Munro, 2000; Jiang et al., 2021; Takahashi et al., 2002). In *Drosophila*, we previously determined that the C-terminal 356 amino acids of PLP (PLP^2539-2895^, here on PLP^C-term^), which includes the PACT domain, interacts with six core centriole and centrosome proteins, plus a single self-interaction (Galletta et al., 2016; Galletta et al., 2014). One attractive model is that some interactions are required to anchor PLP^C-term^ to the centriole, while others play downstream roles in recruiting PCM. To test if deletion of R2720 disrupts specific protein-protein interactions, we performed yeast two-hybrid (Y2H) analysis using both PLP^C-term-WT^ and PLP^C-term-ΔR^ against the seven interactors with this region (Galletta et al., 2016). We found that PLP^C-term-ΔR^ maintained interaction with 5 of the 7 interactors (Fig. 7A,B; S12A), including CaM, confirming our IP result. The two interactions completely disrupted in PLP^C-term-ΔR^ were the intramolecular interaction with PLP^584-1376^ and an interaction with the C-terminus of Asterless^626-994^ (Fig. 7A, B; S12A interactions #3 and #7). We are less confident in the intramolecular interaction (PLP^C-term^ - PLP^584-1376^) because the wild-type interaction was quite weak (Figure 7B; S12A). We therefore hypothesize that the major disruption of PLP^ΔR^ is the loss of interaction with Asterless (Asl), a bridge protein that is anchored to the centriole and important for recruiting PCM (Blachon et al., 2008; Conduit et al., 2014; Varadarajan and Rusan, 2018; Varmark et al., 2007).

**Figure 7:**
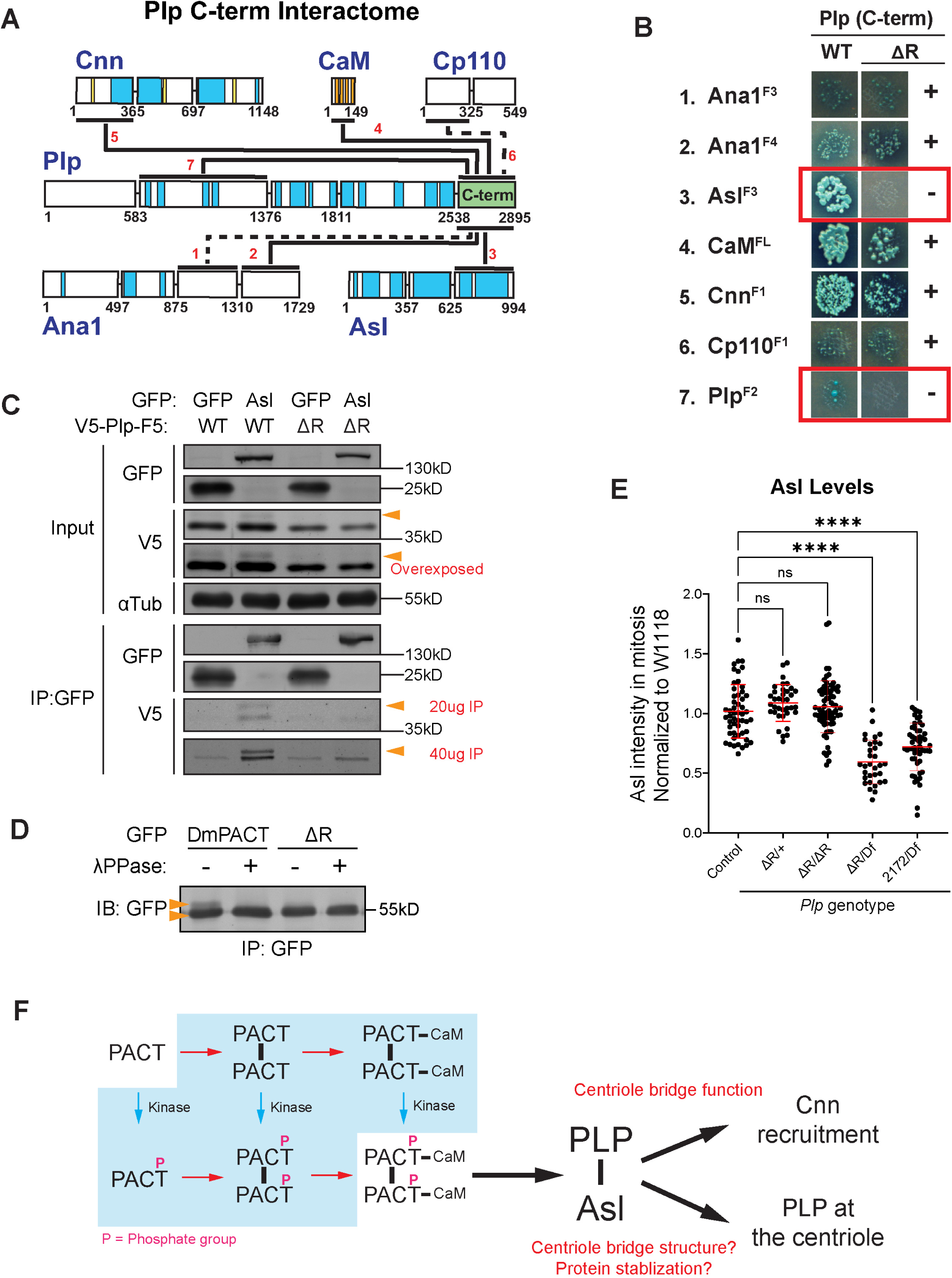
*Plp^ΔR^* does not interact with Asterless. **A.** Map of interactome of the C-terminus (green) of PLP from yeast-two hybrid (Y2H; (Galletta et al., 2016). Solid lines represent scoring 2 or 3. Dashed lines scored 1 (Methods). **B.** Y2H experiment showing interactions of centriole and PCM proteins with PLP(C-term)^WT^ or PLP(C-term)^ΔR^; the PACT domain resides in this fragment. **C.** V5-PLP(c-term) co-IPs with Asl-GFP but not with PLP(C-term)^ΔR^. Note, tight doublet of wild-type PLP(C-term), due to phosphorylation producing the slower migrating species (orange arrow). PLP(C-term)^ΔR^ is not phosphorylated and, thus, the slower migrating band is not observed. **D.** Immunoprecipitated GFP-DmPACT and GFP-DmPACTΔR, mock or lambda phophatase (λPPase) treated. Arrowheads highlight the two species observed. **E.** Asterless intensity at metaphase centrosomes normalized to *w^1118^* controls. Numbers of samples measured are in Supplementary File 1. n.s.= not significant. **** p ≤ 0.0001. **F.** Diagram synthesizing the molecular information gained from this study. The model hypothesizes a molecular pathway that leads from the most basic species (PACT) to a final species of phosphorylated PACT dimer bound to CaM. The exact path taken in cells in unknown (Blue Box), but we presume that a monomer must precede a dimer, and a dimer precedes CaM binding (from this work and Galletta et al., 2014). The final species then promotes PLP interaction with Asl, which is important for setting PLP levels at the centriole, via establishing proper protein levels and / or binding to the centriole. This species at the centriole is also critical for regulating Cnn recruitment.

Consistent with the Y2H data, we found that V5- PLP^C-term^ co-IPed with full-length Asl-GFP (Fig 7C, lanes 1 and 2; S12B,D). Interestingly, immunoblots revealed that PLP^C-term^ appears as a tight doublet, possibly due to its phosphorylation (Fig. 7C, arrowheads, S12B); we observed a similar doublet when either DmPACT or HsPACT are expressed in S2 cells (Fig. 6D,F; orange arrows; S10A, D). Notably, in the Asl-WT lane, the input shows a majority of the V5-PLP^C-term^ is the faster mobility band. In contrast the IP of Asl shows nearly equal amount of the slower and faster bands. This indicates that Asl strongly favors interaction with the slower mobility band (Fig. 7C, IP-V5 vs input-V5), but is still capable of interacting at some level with the high mobility band. To test whether this electrophoretic shift was due to phosphorylation, we treated immunoprecipitates of PLP^C-term-WT^ and PLP^C-term-ΔR^ with Lambda Phosphatase and examined its electrophoretic mobility on SDS-PAGE (Fig. 7D). We found that indeed a subpopulation of PLP^C-term-WT^ was phosphorylated (Phos-PLP^C-term-WT^), appearing as a slower migrating species that disappeared after λ-phosphatase-treatment. Most importantly, PLP^C-term-ΔR^ does not associate with Asl (Fig. 7C, lanes 3 and 4; S12B, D, the higher mobility band in GFP::Asl IP is comparable to control GFP IP). As a critical control, deletion of R2720 in DmPACT or K3154 in HsPACT had no effect on its association with the PCM component Cnn (Fig. S12C), demonstrating the specificity of lost Asl interaction.

Taken together, we hypothesize that Asl binds phosphorylated-PLP with high affinity, and that loss of R2720 blocks PLP phosphorylation, and in turn, Asl binding. The complete mechanistic relationship between phosphorylation, dimerization, interaction with CaM, and interaction with Asl remains to be full determined.

Finally, to test if the reduction in PCM observed in *Plp^ΔR^* was a results of losing a functional interaction between PLP and Asl, or simply because Asl levels are reduced at the centriole, we measured Asl at metaphase centrosomes in wing discs. We found that Asl levels are not altered in *Plp^ΔR/ΔR^* (Fig.7D), although they are reduced in *Plp^ΔR/Df^* and *Plp^2172/Df^*, which suggests that another region of PLP, outside the PACT domain is partially responsible for Asl recruitment. Our previous work did identify a second region of interaction between PLP and Asl (Galletta et al., 2016). Given that the *Plp^ΔR/ΔR^* genotype does not alter Asl levels at the centriole, yet does lead to the reduction in Cnn levels (Fig. 5C), it suggests that the major role for the interaction of PACT and Asl is not in recruiting Asl to the centriole. Instead, we favor a model where the PLP-Asl interaction, which relies on the presence of R2720, is required for efficient recruitment of Cnn to the centrosome.

## Discussion

### *Drosophila* as a model of human disease

One major advantage of studying human disease in *Drosophila* is the ability to assess the effects of an allele across multiple cell types and tissues within a whole organism. Studies in *Drosophila* also occur on a rapid timescale and at significantly lower cost than in vertebrate models. In this study, we modeled a rare, disease-associated variant allele in human *Pericentrin* (p.Lys3154del) using the *Drosophila* ortholog *Plp*. We used CRISPR to generate the orthologous mutation, termed *Plp^ΔR^*, and performed organismal, tissue, cellular, and molecular analyses to characterize its effects at multiple levels. These experiments uncovered phenotypes that parallel some of the human disease presentations including reduced organ size and sensory defects and provides insight into the potential pathogenic mechanisms of this variant.

#### *Plp^ΔR^* is a hypomorphic, separation-of-function mutation with tissue and cell specific effects

It is known that *Plp* null animals die soon after eclosion as a result of severe sensory defects (Martinez-Campos et al., 2004). In contrast, *Plp*^ΔR^ homozygous animals survive as adults, similar to controls, indicating that this allele is likely a hypomorph. Interestingly, we observe that the *Plp*^ΔR^ allele has effects in some tissues, while other tissues are not affected. We see significant effects on wing and head sizes, but the adult brain size is not affected. While there are possible technical explanations for this difference, like fixation and preparation for imaging masking a brain size difference, it is also possible it reflects compensatory mechanisms unique to *Drosophila* neurodevelopment, or that other parts of the head, such as the ocular system might be affected, which would in-turn result in a smaller head size. Spermatogenesis also appears grossly normal, with *Plp*^ΔR^ flies having seminal vesicles full of sperm and the homozygous animals thriving as a stock in the lab. This is particularly interesting as the *Plp* null mutant has substantial defects in sperm development (Galletta et al., 2014; Galletta et al., 2020; Martinez-Campos et al., 2004); (Roque et al., 2018). When we examined sensory function, *Plp*^ΔR^ exhibited an intermediate phenotype between wild-type and null animals. The *Plp*^ΔR^ mutants display impaired mechanosensation and climbing response, consistent with sensory-related phenotypes seen in human patients with MOPD II or other related centrosome-associated microcephaly syndromes (Chen et al., 2019; Endoh-Yamagami et al., 2010; Guirgis et al., 2001; Muhlhans et al., 2011). While previous studies reveal that *Plp* null flies have severe organizational defects in sensory structures and die soon after eclosion as a result of uncoordination (Martinez-Campos et al., 2004), the sensory defects in *Plp*^ΔR^ are present but less severe. To understand the origin of the defects we observed, future experiments examining the JO and other sensory neurons with additional markers, especially those of the cilia, will be needed. Furthermore, our data does not eliminate the possibility that the sensory defects arise from an issue related to the larger cellular context required for sensory function, for example the state of the associated accessory cells of sensory structures and proper neuronal connections.

Together, our modeling indicates that the *Plp*^ΔR^ allele has tissue specific effects. This observation aligns with our previous separation-of-function study involving mutations in the PACT domain (Galletta et al., 2014) and suggests that these tissue-specific roles depend on distinct protein regions of PLP. Such specificity may have implications for human disease, potentially explaining some of the phenotypic variation observed among individuals carrying different variant alleles of the same gene.

#### *Plp*^ΔR^ has multiple effects on development of the wing epithelia

The phenotype we observed that is most consistent with the small stature of MOPD II patients is the reduced wing size, so we examined its development in more detail. While our analysis of the consequences of *Plp*^ΔR^ on the development of the wing epithelia only scratches the surface, our initial finding suggests that the small size may arise from a combination of factors and not from a single issue. We discovered *Plp*^ΔR^ wing discs had an decrease in the number of cells actively dividing and an increase in the number of cells dying. Both could be a contributing factor to the smaller wings of *Plp*^ΔR^ flies, although it is unclear if these phenotypes are independent or interdependent. Furthermore, it is also possible that smaller cell size could contribute to the smaller tissue. Surprisingly, despite the increase in cell death, we did not identify any gross mitotic defects, such as lagging chromosomes or chromosome segregation errors in our fixed and live analyses. It remains possible that errors such as UFBs are present, but are undetectable by light microscopy. Thus, an interesting future research direction is to investigate a link between defective PCNT and the DNA damage response that could then trigger cell death. This phenomena is not limited to wing discs, as we detect cell death in other imaginal discs. This suggests that cell death might be a broad, and perhaps systemic, issue in *Plp*^ΔR^ flies and could provide a hint as to a developmental issue that contributes to MOPD II pathophysiology.

We also observed a faster progression through mitosis (NEB to AO), which could explain part of the decreased mitotic index. It is possible that cell death is a downstream consequence of the accelerated mitosis that leads to errors in chromosome segregation, which then triggers apoptosis. The shortened mitosis could implicate a defective spindle assembly checkpoint (SAC) or premature activation of the anaphase-promoting complex (APC) (Musacchio, 2015). There are phenotypic similarities between *Plp*^ΔR^ and SAC mutants. In the fly, both accelerated mitosis and increased cell death are seen in the brain of mutants in the SAC protein *bub1* (Basu et al., 1999). In mammals, dominant negative Bub1 causes an accelerated exit from mitosis (Taylor and McKeon, 1997) and depletion of an Mad2, another SAC protein, caused cells to enter anaphase prematurely (Gorbsky et al., 1998). Finally, there are many similarities between patients with variants in *PCNT* and SAC, including at the cellular level. Mutations in SAC proteins, including BUBR1, BUB3 and TRIP13, are linked to Mosaic Variegated Aneuploidy (MVA) syndrome, which causes a variety of phenotypes that overlap with MOPD II, including microcephaly, developmental delay, growth limitation, and short stature (de Voer et al., 2013; de Wolf et al., 2021; Hanks et al., 2004; Matsuura et al., 2006; Snape et al., 2011; Yost et al., 2017). Cells from patients with MVA have aneuploidy in a wide range of cell types and have premature sister chromatid separation (PCS) (Hanks et al., 2004; Matsuura et al., 2006). Mitotic spreads of MOPD II (*PCNT* deficient) patient peripheral lymphocytes also have “low-level” MVA and PCS, which lead Rauch et al. to propose a link between PCNT/MOPDII and SAC/MVA (Rauch et al., 2008). Since then, MVA syndrome has been linked to other centrosome proteins, Cep57 and CCDC84 (CENTAC) (de Voer et al., 2013; de Wolf et al., 2021; Snape et al., 2011; Yost et al., 2017). In total, this data suggests a link between the centrosome, PLP/PCNT specifically, and SAC that merits future investigation and could underly the shortened mitosis we observed.

#### *Plp*^ΔR^ provides insights into the PACT domain and its function in PLP

The combination of our fly model, *in silico* modeling, and *in vitro* studies provide important insight into how the Pericentrin PACT domain functions at the centriole (Fig. 7F). Our *in silico* and co-IP data indicate that the PACT domain forms a multimer, likely a dimer, that facilitates binding to CaM. PLP^ΔR^ disrupts coiled-coil formation within the PACT domain, and reduces, although does not eliminate, the ability of PACT to interact with CaM. This is consistent with our previous observations that missense mutations in CBD1 did not disrupt PLP C-terminal binding to CaM (Galletta et al., 2014), as it could still bind PACT via CBD2. However, deletion of the entire CBD1 domain had a much more potent effect as it eliminated CaM binding completely, likely due to disruption in PACT’s ability to dimerize. Collectively, we conclude that CBD1 of PACT is critical for dimerization, and dimerization enhances CaM binding to CBD2, possibly due to a dimerization-induced conformational change in PACT that places CBD2 in an optimal position to bind CaM, all of which was supported by AlphaFold3 predictions.

We show that PLP^ΔR^ prevents phosphorylation of PACT and significantly reduces the interaction between the C-terminus of PLP and the bridge protein Asterless/Asl. We propose that the loss of the PLP-Asl interaction has two downstream consequences: first, it leads to a reduction in the amount of PLP at the centriole, although this could be a result of a reduction in total PLP levels. Second, the loss of the PLP-Asl interaction reduces Centrosomin/Cnn levels at the centrosome. These results lead us to hypothesize that the PLP-Asl interaction is important within the bridge zone of the centriole to recruit and build a robust PCM. (Fig. 7F). The precise interplay between PACT dimerization, CaM binding, and PACT phosphorylation on the PLP-Asl interaction is not fully understood at this point (Fig. 7F, blue box showing molecular species). This important future work will require additional extensive molecular dissection.

In summary, modeling the human MOPD II patient variant, *PCNT*^ΔK3154^, in *Drosophila* revealed how a single amino acid deletion affects biological processes from the molecular level to the organismal level. We identified several phenotypes that closely model MOPD II symptoms including tissue growth defects and sensory impairments. At the cellular, organelle, and molecular levels, we have identified that PLP^ΔR^ alters mitotic timing, increases apoptosis, impairs centrosomal protein recruitment, and disrupts key protein-protein interactions, providing new insights into the potential etiology of MOPD II in patients with this unique *PCNT* variant. With PCNT being a large protein, able to interact with many proteins, we hypothesize that other MOPD II variants from patients might disrupt other critical protein interactions. The loss of different interactions could result in similar loss of function phenotypes in some cellular contexts, but different phenotypes in others. This could provide some explanation for the variation in clinical presentation seen across MOPDII patients. Future directions of this work will follow-up on the MOPD II p.Lys3154del variant with the exiting prospect of designing or screening for interventions that would rescue the PACT-Asl interaction.

## Materials and Methods

### Fly stocks

*D. melanogaster* were maintained on Bloomington Recipe Fly Food (Lab Express, Ann Arbor, MI). Crosses were performed at 25°C unless otherwise described. All control (wild-type) flies used in this study are *yw* or *w1118*. The following strains were used: *Plp^2172^* (Spradling et al., 1999), *Df(3L)Brd15* (Bloomington Drosophila Stock Center, Bloomington, IN, USA, #5354), H2A::RFP (*His2AvD::mRFP)* (Pandey et al., 2005), ubi-tubulin:GFP (Gift from Tomer Avidor-Reiss Lab), UAS-Ana1::tdtomato (Blachon et al., 2009), ubi-Plp::GFP (Galletta et al., 2014) and BAC-Plp (transgenic line carrying a BAC that duplicates the region around *Plp*; Bloomington Drosophila Stock Center, #90069).

### Fly cell culture

*Drosophila* S2 cells (Invitrogen) were cultured in Sf-900II SFM media (Life Technologies) supplemented with penicillin/streptomycin. DNA plasmid was transfected into S2 cells by nucleofection. Briefly, ∼5 × 10^6^ cells were pelleted by centrifugation. Cell pellet was resuspended in 100 µl of transfection solution (5 mM KCl, 15 mM MgCl2, 120 mM sodium phosphate, and 50 mM D-mannitol, pH 7.2) containing 2 µg of purified plasmid, transferred to a cuvette (2 mm gap size), and then electroporated using a Nucleofector 2b (Lonza), program G-030. Transfected cells were diluted immediately with 0.4 mL SF-900 II medium and plated in a 6-well cell-culture plate containing 1 mL of fresh media. Cells were allowed 24 hours to recover before additional handling. Expression of the construct was induced by the addition of 0.25 mM CuSO4 to the culture medium. Transfected cells were used 24 hours after induction.

### Plasmids and molecular cloning

The CRISPR repair template was cloned into pUC backbone with Gibson assembly using primers:

PLP gDNA Forward: GTACTTCGCGAATGCGTCGAGATACCAATGACCGACGACGAGAACTTCACTGGCG AGCGG

PLP gDNA Reverse: CTCGTCGGTCCCGGCATCCGATCCAATAGGATGATGCCGCGCATGCGCTCTTTTT GGTTT

Site directed mutagenesis was performed to remove R2720 from the repair template using primers:

PLP ΔR2720 mutagenesis Forward: GCTGGTCTACCAGAAGTATTTGAAGCTCACAC

PLP ΔR2720 mutagenesis Reverse:

GTGTGAGCTTCAAATACTTCTGGTAGACCAGC.

Site directed mutagenesis was performed to silently remove a StuI enzyme cleavage site ∼300 bp upstream of R2720 from the repair template using primers:

PLP StuI mutagenesis Forward: GCCAAACTAGCTGAGGCCTTAGCTCAGGC PLP StuI mutagenesis Reverse: GCCTGAGCTAAGGCCTCAGCTAGTTTGGC

Site directed mutagenesis was performed to remove the PAM of the gRNA site from the repair template using primers:

PLP gRNA PAM mutagenesis Forward: CACTGGAAGGTTACCAAGCCAGTGAGCAATTGG

PLP gRNA PAM mutagenesis Reverse: CCAATTGCTCACTGGCTTGGTAACCTTCCAGTG

Primers for cloning the PLP ΔR2720 gRNA into pU6-BbsI-chiRNA vector (Gratz et al., 2013) were:

PLP ΔR2720 gRNA Forward: CTTCGCTCACACTGGAAGGTTACC

PLP ΔR2720 gRNA Reverse: AAACGGTAACCTTCCAGTGTGAGC

Lentiviral mNG-HsPACT (PCNT aa. 3102-3336) plasmid was ordered from Twist Biosciences. mNG-HsPACT-ΔK3154 was generated through PCR mutagenesis of mNG-HsPACT with primers:

ΔK3154-F: gctctgatttatcaaaagtatcttttgctgttgattgg

ΔK3154-R: ccaatcaacagcaaaagatacttttgataaatcagagc

### Generation of CRISPR flies

*PlpΔR2720* was generated by homologous recombination using CRISPR (Gratz et al., 2013). (Fig. 1F). y[1] M(vas-Cas9)ZH-2A w[1118]/FM7c (Bloomington Drosophila Stock Center, #51323) flies were crossed to PLP::mNeon CRISPR line (Galletta et al., 2020). The progeny were injected with pU6-Bbsl-chiRNA containing the PlpΔR2720 gRNA and the repair template (BestGene; Chino Hills, CA). Recombinants were identified by screening PCR products amplified from potential recombinants for loss of digestion by StuI. Counter-screening found this recombinant was untagged and did not contain mNeon.

### Brain volume measurements

Adult female flies aged 4–6 days were collected from the respective genotypes cultured at either 25 °C or 15 °C. Flies were anesthetized, and their heads were separated using dissection forceps, then placed in 4% formaldehyde solution for 15–20 minutes at room temperature. Following this, fly brains were dissected and further fixed for an additional 20 minutes in 4% formaldehyde at room temperature. After fixation, the solution was removed and the dissected brains were incubated in 1× PBS for 30 minutes to ensure removal of any residual fixative. The brains were then incubated in blocking buffer (0.1% PBST with 0.5% BSA) for 1 hour at room temperature, followed by overnight (O/N) incubation with primary antibody at 4 °C. The next day, samples were washed three times with 0.1% PBST and incubated with the appropriate secondary antibody for 2 hours at room temperature, followed by three washes with 0.1% PBST. Samples were then stained with nuclear stain (NuclearMask, Invitrogen H10325, 1:2000 dilution) for 15 minutes, followed by PBST washes. Phalloidin-Atto-647 (Sigma 65906, 1:1000) was then added for 15 minutes, followed by a PBST wash. Finally, samples were washed in 1× PBS to remove any residual Triton. Samples were mounted on glass slides using 0.12 µm depth Secure-Seal Imaging Spacers (Grace Bio-Labs 654008) in Aqua-Poly/Mount (Polysciences 18606-5). Slides were cured at room temperature overnight before imaging. The phalloidin channel was used to perform volumetric analysis across different genotypes. Full Z-stacks were imported into *Imaris* 10.2.0 version software, 3D volumes were rendered using the surpass volume rendering module. Volumetric measurements in µm^3^ were obtained from the statistics tab.

### Bang assay

Bang assays were performed using a previously described modified protocol (Inagaki et al., 2010). The apparatus was 3D printed and consists of an upper and lower frame with five circular indentations into which plastic fly vials (25 mm diameter x 95 mm height) are inserted. The openings of the vials in each frame are faced towards each other and connected by a third frame containing a manual sliding gate mechanism, allowing flies to move freely between the upper and lower vials when open. In brief, 15-30 male flies were collected one day after eclosion under ice anesthesia and transferred to a fresh culture vial. Flies were placed in a dark room for 1-3 hours to recover from anesthesia and acclimate to darkness. To perform behavioral assay, flies were transferred to the lower vial of the apparatus with the sliding gate closed. For each trial, the apparatus was forcefully banged down onto a rubber mat three times to agitate flies and induce strong negative gravitaxis. The apparatus allows for five genotypes to be tested simultaneously to control for variability in agitation force. Immediately after agitation, the sliding gate was opened for 1 min to allow flies to climb from the lower to upper vial. Afterwards, the sliding gate was closed and the number of flies that had climbed the approximately 95 mm into the upper vial was recorded. Flies were given a 5 min recovery period after each trial and repeated for five trials. As a negative control, wild-type flies were collected one day prior to experiment, anesthetized on ice, and aristae in the third antennal segment were removed using fine forceps or iridectomy scissors, referred to herein as ablation. Flies were transferred to a fresh culture vial containing a moistened Kimwipe tent. The second antennal segments containing Johnston’s organs were carefully unperturbed to prevent a compensated gravity response. All experiments were performed between 70-73°C and 30-32% ambient humidity. To determine if the defects seen in this assay were a result of climbing issues, and not solely a result of defects in gravity sensation, these experiments were repeated with a light above the apparatus to induce phototaxis, which should be independent of gravity sensation.

### Mechanosensation assay

Mechanosensation assays were performed using a previously described, modified protocol (Murphy et al., 2015). In brief, flies were collected on the day of eclosion, anesthetized on ice, and decapitated with iridectomy scissors. Decapitation is necessary to prevent adults from flying away when stimulated. Headless flies were placed in a closed, moist chamber and allowed approximately 10-20 hours of recovery. After the recovery period, only flies that right themselves when perturbed were used for further testing. All experiments were performed within 24 hours of decapitation. On the testing day, flies were visualized under a dissecting microscope and a grooming reflex was elicited by deflecting either the notopleural or postalar bristles towards the fly body with a stiff paintbrush or fine forceps. The leg grooming response was given a score of one for flies that lifted their leg in response to bristle stimulation and a score of zero to flies that did not move their legs. For each fly, bristles were stimulated up to three times per trial with five total trials spaced 2 min apart.

### Immunofluorescence

Imaginal discs and testes were dissected, from developmentally staged third-instar larvae or adult males respectively, in Schneider’s medium (ThermoFisher Scientific, Waltham, MA, USA) with antibiotic-antimycotic (ThermoFisher Scientific, Waltham, MA, USA), and fixed in 9% formaldehyde in PBS at room temperature for 20 min. Fixed samples were washed 3 times in PBS + 0.3% Triton X-100 (PBST) and blocked for 2-4 hours in PBST + 5% normal goat serum (NGS). Samples were incubated in primary antibodies in PBST + 5% NGS overnight at 4°C. After 3 x 10 min washes in PBST, samples were incubated for 2 hours at room temperature with secondary antibodies. Samples were finally washed 3 x 10 min in PBST and mounted in AquaPolymount (Polyscience, Inc., Warrington, PA, USA) underneath a #1.5 coverslip. For Johnstons Organ’s antennae were dissected from pupae ∼60 – 70 hours after pupariation. Fixation, staining and mounting were as for other tissues. Primary antibodies used were rabbit anti-PLP (1:5000) (Rogers et al., 2008), guinea pig anti-Asterless (1:10,000; gift from G. Rogers, University of Arizona Cancer Center, University of Arizona, Tucson, AZ, USA), rabbit anti-Centrosomin (1:10,000; gift from T. Megraw, Florida State University, Tallahassee, FL; USA), rabbit anti-phospho-histone H3 (1:1000; MilliporeSigma, Burlington, MA, USA), rabbit anti-cleaved DCP-1 (1:200; Cell Signaling Technology, Danvers, MA, USD), mouse anti- γH2Av (1:100; Developmental Studies Hybridoma Bank, UNC93-5.2.1-s) and rat anti-Deadpan (1:100; Abcam, Cambridge, UK). Secondary antibodies labeled with Alexa Fluor 488, 568, or 647 were used at 1:1000 (ThermoFisher Scientific). Phalloidin and DAPI (1:1000; ThermoFisher Scientific) was added to secondary antibodies.

### Microscopy

Confocal images were collected using a Nikon Eclipse Ti2 (Nikon Instruments, Melville, NY, USA) with a Yokogawa CSU-W1 spinning disk confocal head (Yokogawa, Life Science, Sugar Land, TX, USA) equipped with a Prime BSI cMOS camera (Teledyne Photometrics, Tucson, AZ, USA) and Nikon Elements software (Nikon Instruments). Adult wings and heads were imaged with a 4X air objective. Wing discs were imaged with a 100x TIRF/1.49 NA oil immersion objective for centrosome-specific measurements and a 40x/1.30 NA oil objective for mitotic index measurements. Johnston’s organs were imaged with a 100x TIRF/1.49 NA oil objective on a Nikon Eclipse Ti (Nikon Instruments) with a Yokogawa CSU-X1 spinning disk confocal head (Yokogawa, Life Science) equipped with an Kinetix22 camera (Teledyne Photometrics, Tucson, AZ) and Nikon Elements software (Nikon Instruments). For adult brain volumetric analysis samples were imaged on a Zeiss LSM 880 confocal microscope using a 20×/0.8 NA objective with Zen software. For wing disc measurements, Z-stacks encompassing the entire tissue volume were collected by acquiring optical slices spaced by 200 nm; these data are presented as maximum intensity projections. 405, 488, 561, and 641 nm laser lines were used on all systems. All images were analyzed and processed using Fiji (Image J, National Institutes of Health, Bethesda, MD, USA) unless noted.

S2 cell imaging was performed using a CSU-W1 SoRa Yokogawa spinning-disk field scanning confocal system assembled on an ECLIPSE Ti2 inverted microscope (Nikon) with a Kinetix sCMOS 10MP camera (Photometrix) with a 100x (Silicone Plan Apo, NA = 1.35) objective.

### Fluorescence measurements

Centriole protein measurements were performed as previously described (Galletta et al., 2014). Samples and a corresponding control were prepared and imaged on the same day. Fixed wing disc and brain samples were stained with anti-PLP, anti-Asl, and anti-Cnn to label centrioles and DAPI to label metaphase nuclei. Sum projections of the entire Z volume of the centriole were generated, an ROI was drawn around the centriole using the asterless label for reference, and the total integrated density was measured. An identically sized ROI adjacent to the centriole was measured for background subtraction. All fluorescence measurements were normalized to the mean value of the *yw* or *w1118* control on a given day. The Centrosomin signal was carefully outlined for each centriole and the circularity shape descriptor was measured.

Wing disc mitotic index measurements were performed as previously described (Handke et al., 2014). All samples were prepared and imaged on the same day. Fixed wing disc samples were stained with anti-phospho-histone H3 (PH3) to label cells undergoing mitosis with DAPI and phalloidin to delineate all individual cells. A first focal plane of the wing discs was acquired at the peripodial membrane. Additional optical slices with 200 nm spacing were acquired in the direction towards the basal side of the disc epithelium. Maximum intensity projects of each z stack were generated using Fiji to produce 2D images. The area defined by the pouch and hinge regions of the wing discs was outlined and a mask was applied in the channel containing PH3-positive nuclei. The area occupied by PH3-positive nuclei was divided by the total area of the combined pouch and hinge region to describe a mitotic index of the tissue. To quantify cell death, the same method was employed as previously described except fixed wing disc samples were stained with anti-cleaved DCP-1 to label cells undergoing apoptosis.

### Wing disc western blots

For analysis of PLP levels by Western blot. 30 wing discs from crawling third instar larvae were dissected in Schneider’s media with antibiotic-antimycotic and homogenized in 50 µLof RIPA buffer (10 mM Tris-HCl, pH 7.5, 140 mM NaCl, 1 mM EDTA, 0.5 mM EGTA, 1% Triton X-100, 0.1% sodium deoxycholate, 0.1% sodium dodecyl sulfate). 10 µL of 6X SDS-sample buffer (350 mM Tris pH 6.8, 30% glycerol, 10% SDS, 600mM DTT, ∼0.05% Bromophenol Blue) was added and samples were incubated for 5 min at 95°C, then samples were stored at -80°C until use. All of each sample was run on a 1.5mm thick 5% SDS-polyacrylamide gel. Samples were then transferred to Polyvinylidene fluoride (PVDF) membrane using Tris-Base Glycine transfer buffer (Novex) with 20% methanol. Blots were blocked in 5% nonfat dry milk diluted in TBST (0.1% Tween 20 diluted in Tris Buffered Saline—50-mM Tris-HCl, pH7.5, 150-mM NaCl) for 30–60 min before incubation with primary antibody - Rabbit anti-PLP (N-terminus, (Rogers et al., 2008); 1:2000) - diluted in block overnight at 4°C. Blots were rinsed 3X, then washed 3 x 10 minutes in TBST. Secondary antibody (anti-Rabbit horseradish peroxidase (HRP) conjugated 1:5000; ThermoFisher) in block for at least 2.5 hours at RT. Blots were rinsed 3X, then washed 3 x 10 minutes in TBST. Detection used SuperSignal West Dura Extended Duration Substrate (Life Technologies) and an Amersham ImageQuant 800 (Cytiva). To assess loading, blots were stripped with Restore Wesern Blot Stripping Biffer (ThermoFisher) and reprobed with anti-α spectrin (1:50; 3A9; Developmental Studies Hybridoma Bank) and anti-mouse-HRP (1:5000; ThermoFisher) as above. For quantification, “total integrated density” of bands were measured in ImageJ and background from a region of the same size, adjacent to the that band was subtracted. Normalization was first to the loading control (α-Spectrin) and then to the control PLP band of that experiment. Mean ± standard deviation of three independent experiments is presented.

### Live imaging

For live imaging of wing discs, crawling third-instar larvae were dissected in Schneider’s medium with antibiotic-antimycotic. Wing discs were mounted in a drop of Schneider’s medium onto a gas-permeable Lumox tissue culture dish (Sarstedt, Newton, NC). Imaging was performed using a Nikon 40×/1.30 NA oil objective on a Nikon Eclipse Ti (Nikon Instruments) with a Yokogawa CSU-X1 spinning disk confocal head (Yokogawa, Life Science) equipped with a Kinetix22 camera (Teledyne Photometrics, Tucson, AZ), 1.5x tube lens, and Nikon Elements software (Nikon Instruments). A Tokai Hit Stage Top Incubator (Tokai Hit USA Inc., Bala Cynwyd, PA) was used to maintain a constant temperature of 25°C throughout imaging. The first focal plane was acquired at the peripodial epithelium. Seven optical slices with 2 µm spacing were acquired every 20s for one hour. The Nikon Perfect Focus System (PFS) was used to continuously monitor the focal plane and make adjustments to keep samples in focus.

### Protein *in silico* Structure Analysis

The coiled-coil regions of DmPACT^WT^ and DmPACT^ΔR^ were predicted by submitting the PACT sequence (AA 2634-2780) of PLP (Gene ID: 3772382) to WaggaWagga, which includes Multicoil and NCOILS predictions (Simm et al., 2021).

Molecular modeling was carried out using the AlphaFold 3 server (Abramson et al., 2024). The DmPACT^WT^ and DmPACT^ΔR^ sequences (AA 2634-2780) and *Drosophila* calmodulin sequence were used for modeling (Gene ID: 26329). All structural figures were created with PyMOL (Version 3.1.1).

### Yeast two-hybrid

PLP interactions were tested using the yeast two-hybrid assay as previously described (Galletta and Rusan, 2015). In brief, cDNA sequences of PLP and PLP-interactor fragments (cloning information listed above) were cloned into pDEST-pGADT7 (Rossignol et al., 2007) and pDEST-PGBKT7-Amp (Galletta et al., 2014) using the Gateway cloning system (Life Technologies) and transformed into Y187 and Y2HGold strains, respectively. The transformants were cultured in either Sd-Leu or Sd-Trp media to select for those carrying the appropriate vector. Strains to be tested for interactions were mated by mixing Y187 and Y2hGold strains in yeast extract + peptone + adenine + dextrose (YPAD) medium overnight at 30°C with shaking in a flat bottom 96-well plate. Diploids were selected by plating on SD-Leu-Trp dropout media (DDO). Diploids were replicated onto test plates: DDO to control for diploid growth and QDOXA (DDO + Aureobasidin A + X-α-Gal) for interaction tests. Interactions were scored from test plates based on the presence of growth and development of blue color on the QDOXA plate on a scale of 0 (no growth) to 3 (robust growth and blue color).

### Immunoprecipitation

Immunoprecipitation (IP) assays were performed using recombinant, purified GFP-Binding Protein (GBP) fused to human Fc domain and bound to magnetic Protein A Dynabeads (ThermoFisher) and then cross-linked with dimethyl pimelimidate. GBP-coupled Dynabeads were stored in PBS, 0.1% Tween 20 at 4°C. Before use, beads were equilibrated in IP buffer (50 mM Tris, pH 7.2, 125 mM NaCl, 1 mM DTT, 0.5% Triton X-100, 1x SigmaFast Protease inhibitors, 0.1 mM <u>PMSF</u>, and 1 μg/mL soybean trypsin inhibitor). Transfected cells were harvested and lysed in IP buffer. Lysate concentration was determined by Bradford assay, and lysates were diluted to 5 mg/mL. Lysates were then clarified by centrifugation at 10,000 x g for 5 min at 4°C. Inputs were made from pre-cleared lysates in Laemmli Buffer. GBP-coated beads were added with lysates and rocked for 30 min at 4°C, followed by four 5-minute washes by resuspending beads in 1 mL IP buffer, and then transferred to a new tube during the final wash, then boiled in 2x Laemmli sample buffer. For the lambda phosphatase treatment, lysates with GBP-coated beads were incubated with lambda phosphatase for 30 min and then boiled in 2x Laemmli sample buffer. Detection by Western Blot utilized anti-V5 (ThermoFisher) and anti-GFP JL-8 (ThermoFisher).

### Statistical analysis

Quantified experiments were performed 2 or 3 times to ensure reproducibility. Data analysis and statistics were performed using Excel (Microsoft) and Prism (GraphPad version 10.4.1) software. For comparisons between two groups, unpaired two-tailed *t-*tests were used, with Welch’s correction when appropriate. One-way ANOVA with Tukey’s correction was used for comparisons between three or more groups. Sample sizes are reported in the figure legends and/or Supplemental File 1. Error bars represent the mean ± SD for all graphs.

## Acknowledgements

We thank the NHLBI light microscopy core for assistance with the Zeiss LSM 880 microscope. This work was supported by the Division of Intramural Research at the National Heart, Lung, and Blood Institute (ZIAHL006126 to NMR), the National Institute of General Medical Sciences (R35GM136265 to GCR), and the National Cancer Institute (P30CA23074 to the UACC).

## DISCLAIMER

This research was supported by the Intramural Research Program of the National Institutes of Health (NIH). The contributions of the NIH author(s) are considered Works of the United States Government. The findings and conclusions presented in this paper are those of the author(s) and do not necessarily reflect the views of the NIH or the U.S. Department of Health and Human Services.

## DATA AVAILABILITY STATEMENT

All relevant data can be found within the article and its supplementary information. The DOI for all raw data is 10.25444/nhlbi.30657017

## SUPPLEMENTAL MATRIAL

Figure S1: Head size images and measurements.

Figure S2: Wing images and measurements

Figure S3: Phototaxis assay

Figure S4: *Plp^ΔR^* does not show gross defects in Johnston’s Organ

Figure S5: Wing imaginal disc mitosis and apoptosis imaging.

Figure S6: Eye/Antennal and leg imaginal disc mitosis and apoptosis imaging.

Figure S7: PlpΔR affects interphase PLP levels at the centriole and total wing disc levels.

Figure S8: *Plp^ΔR^* impacts PCM recruitment.

Figure S9: AlphaFold3 structural predictions of DmPACT.

Figure S10: *Plp^ΔR^* affects protein-protein interactions.

Figure S11: AlphaFold3 structural predictions of HsPACT.

Figure S12: *Plp^ΔR^* affects interaction with Asl.

Supplemental File 1: Numbers of samples measured for all quantifications.

