## Supplemental Figures for "A MOPD II-associated Pericentrin variant disrupts PACT domain dimerization and pericentriolar material recruitment"

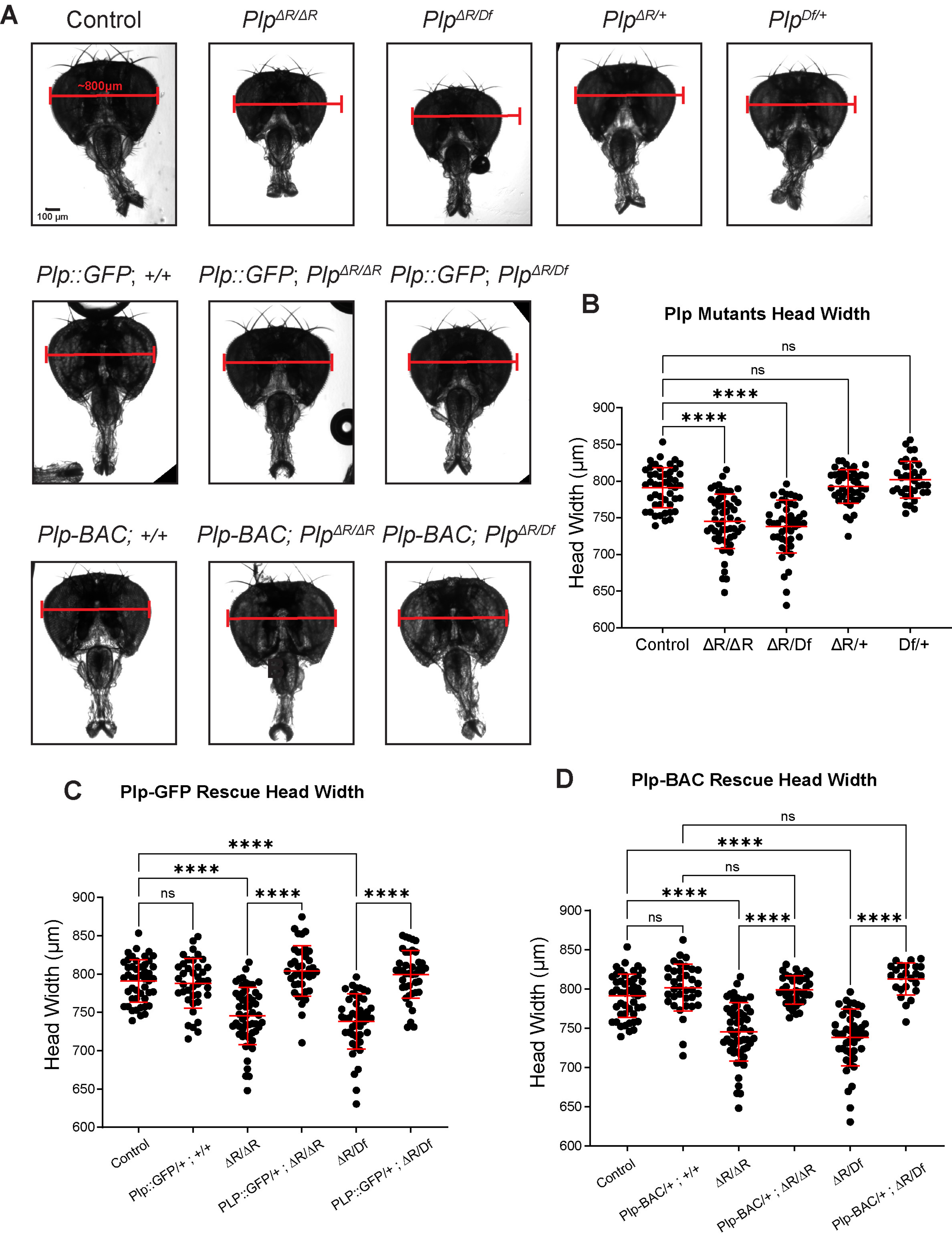


Thomas and Galletta et al., Figure S1

**Figure S1: Head images and measurements**

Some images and measurements have been duplicated from Figure 2 to for ease of comparison. Some measurements are duplicated within Figure S1 for ease of comparison. **A.** Adult heads. 800 μm bars for ease of comparison. **B.** Adult head width measurements, mutants, and heterozygotes. **C.** Adult head width measurements, mutants, ubi-PLP::GFP rescues, and controls. **D.** Adult head width measurements, mutants, BAC-PLP rescues, and controls. Numbers of samples measured are in Supplementary File 1. n.s. = not significant, ****$p \leq 0.0001$.

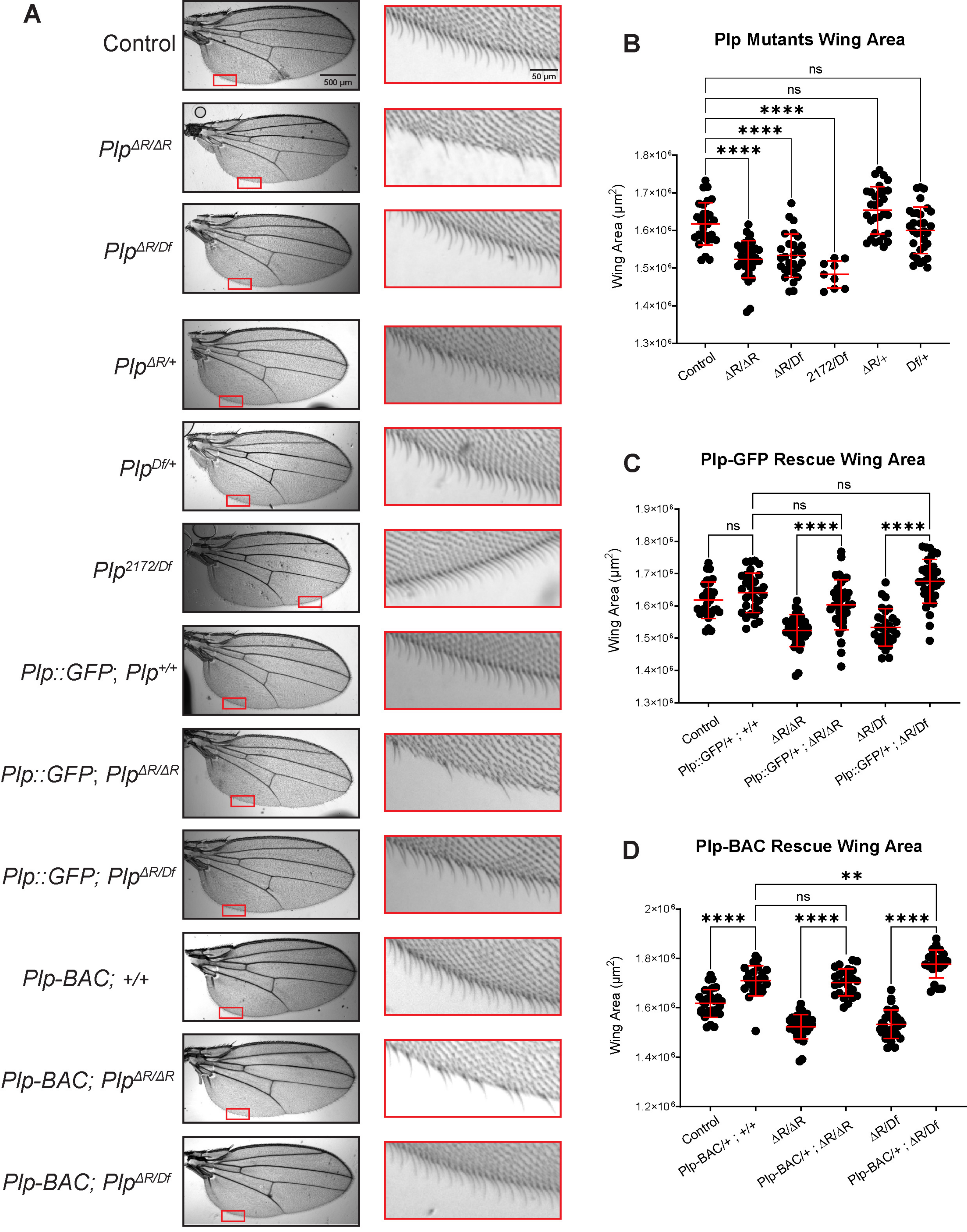


Thomas and Galletta et al., Figure S2

**Figure S2: Wing images and measurements**

Some images and measurements have been duplicated from Figure 2 to for ease of comparison. Some measurements are duplicated within Figure S2 for ease of comparison. **A.** Adult wings. Red insets of wing margin shown at higher magnification, right. **B.** Adult wing area measurements, mutants, and heterozygotes. **C.** Adult wing area measurements, mutants, ubi-PLP::GFP rescues, and controls. **D.** Adult wing area measurements, mutants, BAC-PLP rescues, and controls. Numbers of samples measured are in Supplementary File 1. n.s. = not significant, $**p \leq 0.01$, $****p \leq 0.0001$.

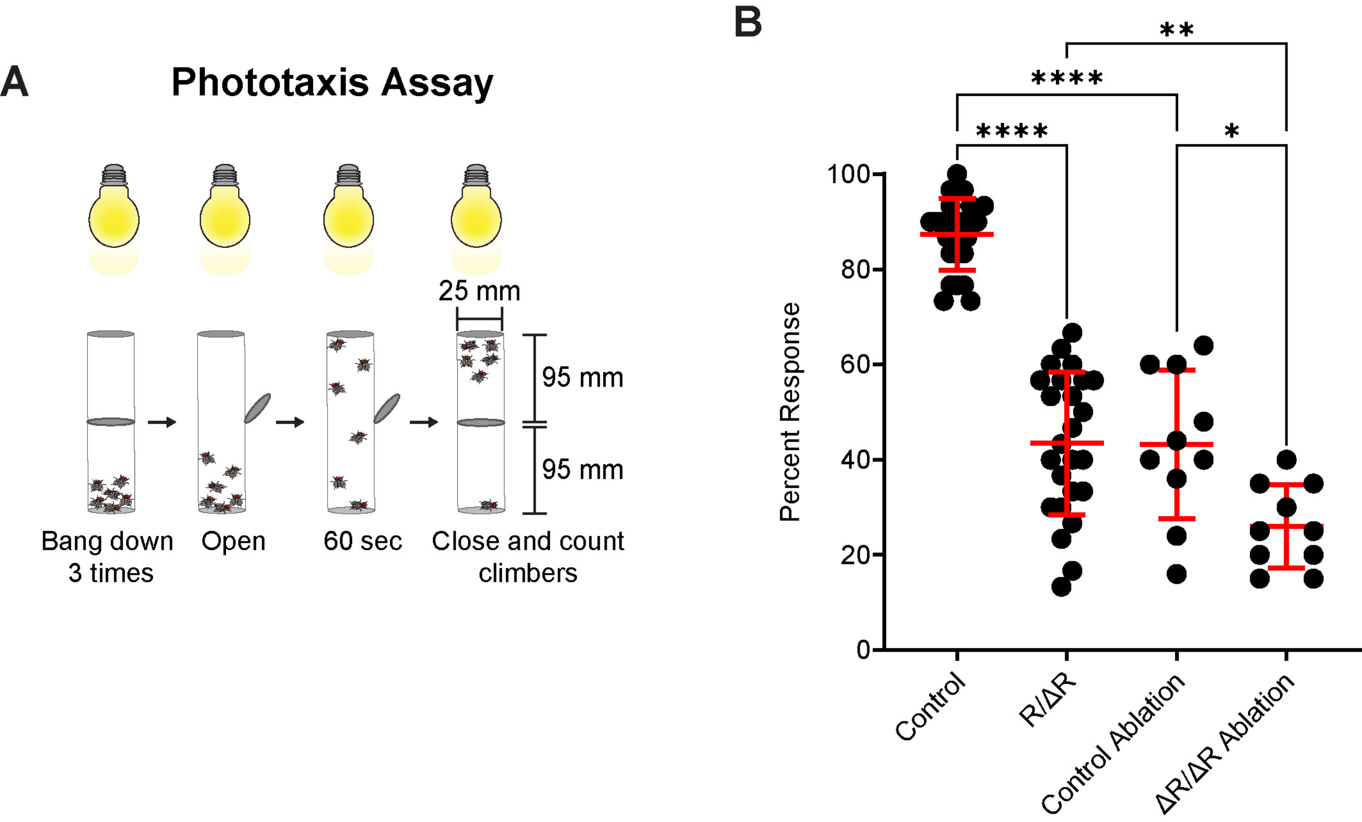


Thomas and Galletta et al., Figure S3

**Figure S3: Phototaxis assay**

**A.** Schematic of phototaxis assay. **B.** The percentage of flies that climbed to upper chamber within 60s is reported. Each point represents the average climbing response from a group of 15-30 flies assayed three times. Numbers of samples measured are in Supplementary File 1. *$p \leq 0.05$, ***$p \leq 0.001$, ****$p \leq 0.0001$.

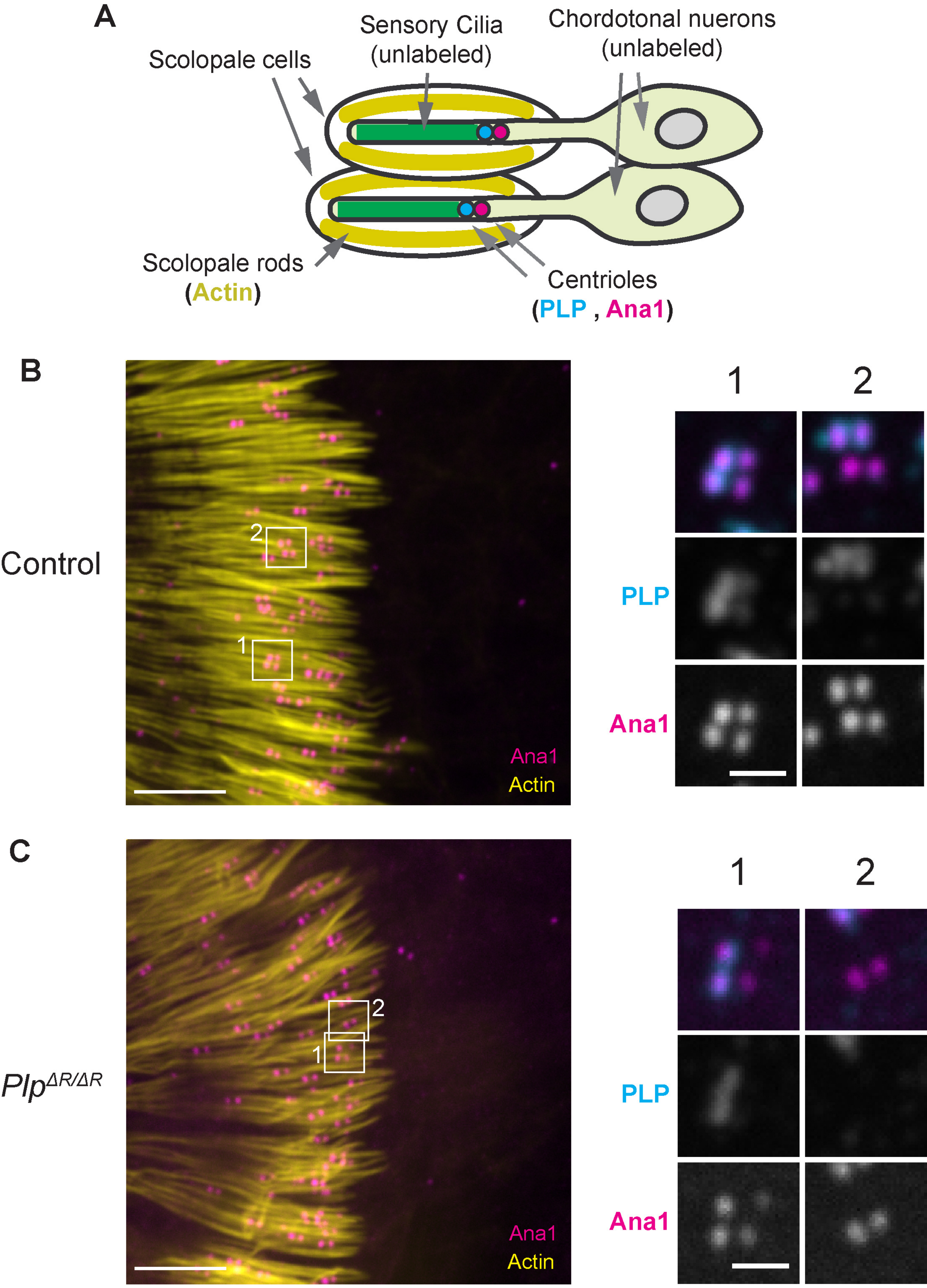


Thomas and Galletta et al., Figure S1

**Figure S4: *plp*$^{\Delta R}$ does not show gross defects in Johnston's Organ**

**A.** Cartoon of the cells of the *Drosophila* Johnson's Organ and the location of the centrioles and cilia. **B-C.** Micrographs of a portion of the Johnston's Organs of **(B)** control and **(C)** *plp*$^{\Delta R}$ antennae. Actin (yellow, phalloidin), Ana1 (magenta, Ana1::tdtomato) and PLP (Cyan, anti-PLP immunofluorescence). The organization of the centrioles of the mutant appears grossly normal relative to the scolopale rods (actin). PLP is present on one centriole of each pair (top right), although not in all cases (bottom right) of both control and ΔR. Images are presented for optimal viewing of each channel in each image and should not be compared for intensity across images. Scale Bars: left – 5 μm, right – 1 μm

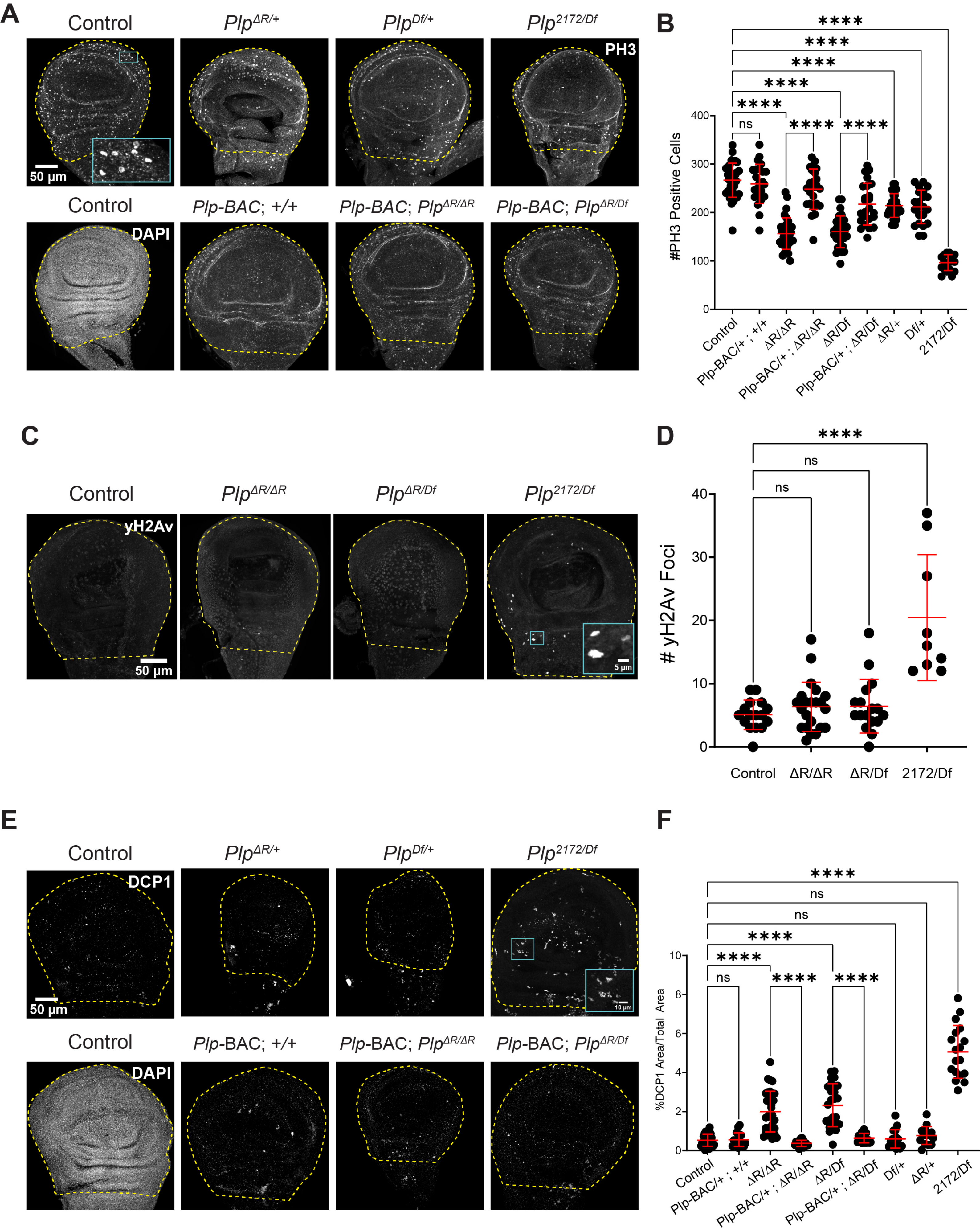


Thomas and Galletta et al., Figure S5

**Figure S5: Wing imaginal disc mitosis and apoptosis imaging.**

Figure is supplement to Figure 4. Some images and measurements have been duplicated from Figure 4 to for ease of comparison. **A.** Wing imaginal discs stained for DNA (Control, DAPI, lower left) and active mitosis (phospho-histone H3 (PH3), all others). The combined wing pouch and hinge region used for analysis are outlined. **B.** Number of PH3-positive nuclei in total wing pouch and hinge area. **C.** Wing imaginal discs stained for double stranded DNA breaks (γH2Av). The combined wing pouch and hinge region used for analysis are outlined. **D**. Number of γH2Av foci in the total wing pouch and hinge area. **E.** Wing imaginal discs stained for DNA (DAPI, Control, lower left) and an apoptosis marker(cleaved *Drosophila* Death caspase-1 (DCP1), all others). The combined wing pouch and hinge region used for analysis are outlined. **F.** Percentage of DCP1-positive area out of total wing pouch and hinge area. Numbers of samples measured are in Supplementary File 1. n.s.= not significant. ****$p \leq 0.0001$.

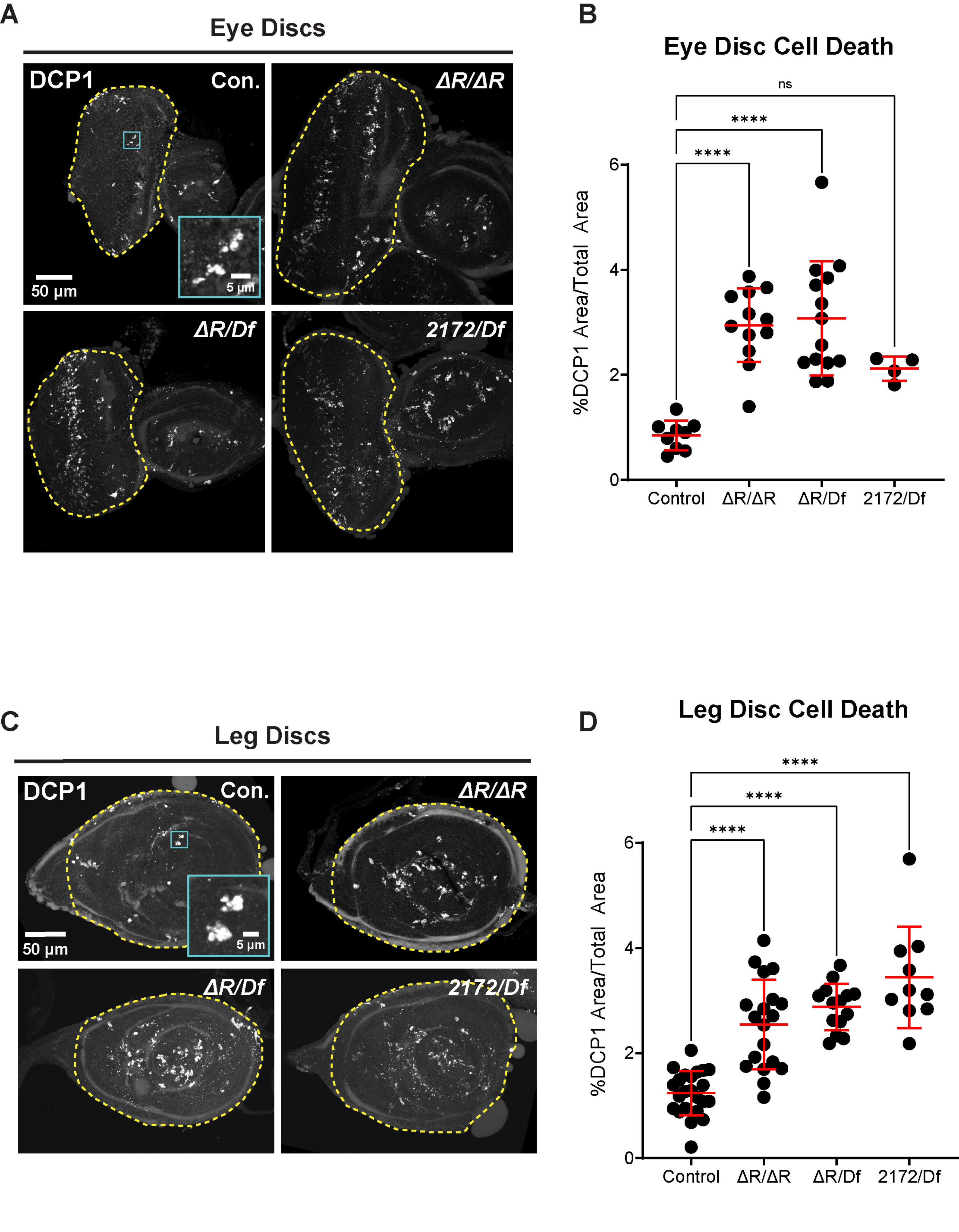


Thomas and Galletta et al., Figure S6

**Figure S6: Eye/Antennal and leg imaginal disc mitosis and apoptosis imaging.** **A.** Wing imaginal discs stained for apoptosis (cleaved DCP1). The eye disc areas used for analysis are outlined. **B.** Percentage of DCP1-positive area out of total eye disc area. **C.** Leg imaginal discs stained for apoptosis (cleaved DCP1).The leg disc areas used for analysis are outlined. **D.** Percentage of DCP1-positive area out of total leg disc area. N-values are in Supplementary File 1. n.s.= not significant. ****p ≤ 0.0001.

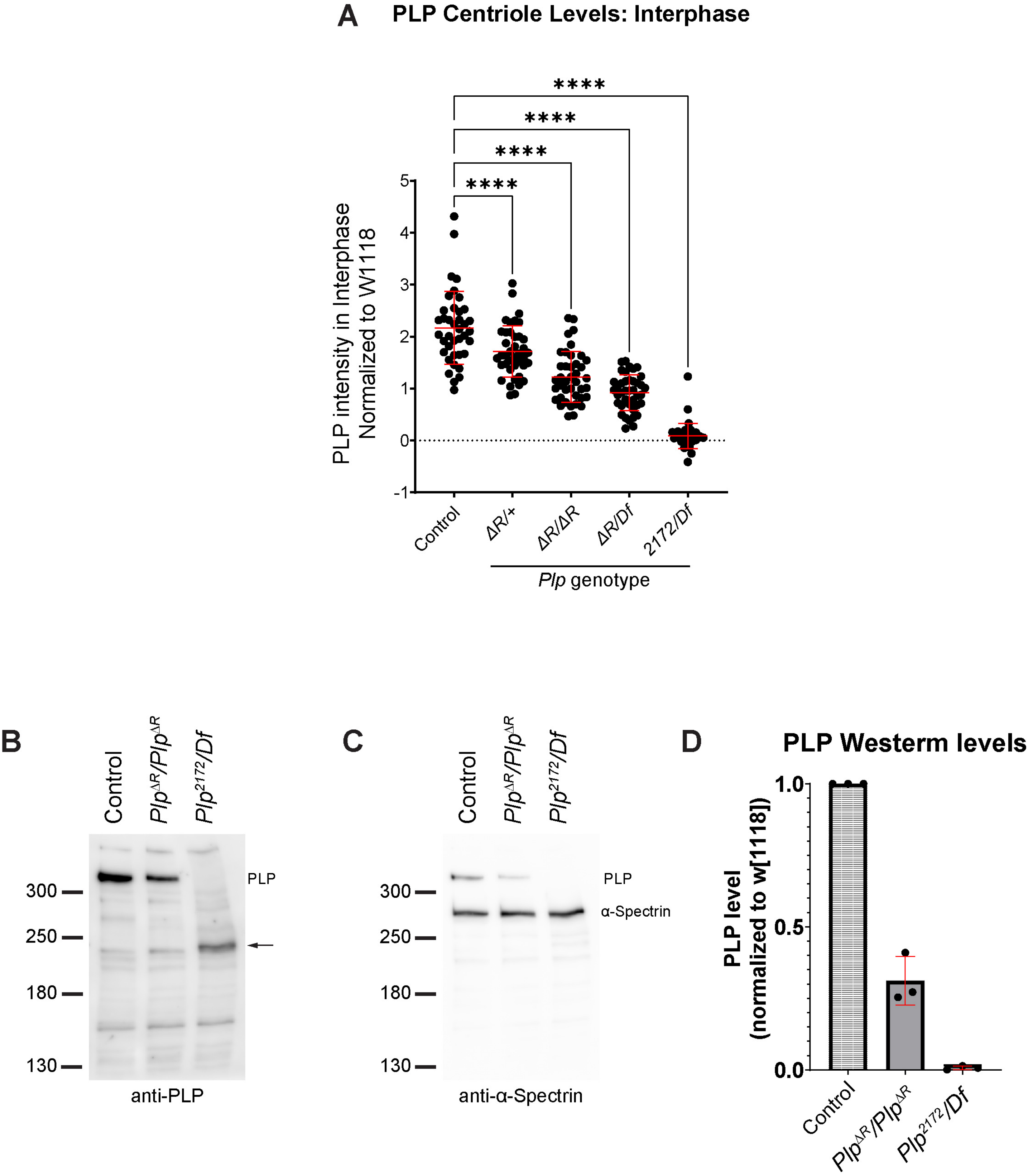


Thomas and Galletta et al., Figure S7

**Figure S7: *Plp$^{\Delta R}$* affects interphase PLP levels at the centriole and total wing disc levels.**

**A.** PLP intensity at interphase centrioles in wing disc cells normalized to *w$^{1118}$* controls at metaphase (Fig. 5C). ****p ≤ 0.0001. Numbers of samples measured are in Supplementary File 1.. **B-D.** Western blots of total extract from 30 wing discs from flies of the indicated genotypes. **B.** anti-PLP Blot. Full length PLP is noted. Arrow indicates band seen only in *Plp$^{\Delta R/Df}$* likely representing a C-terminally truncated product. **C.** Reprobe of blot in B after stripping. anti-α Spectrin as loading control. Note some signal from the anti-PLP blot was not stripped. **D.** Quantification of full length PLP levels in indicated genotypes. Normalized for loading and to control levels. Three independent repeats of dissection, lysis and blotting were performed. Mean ± standard deviation is presented.

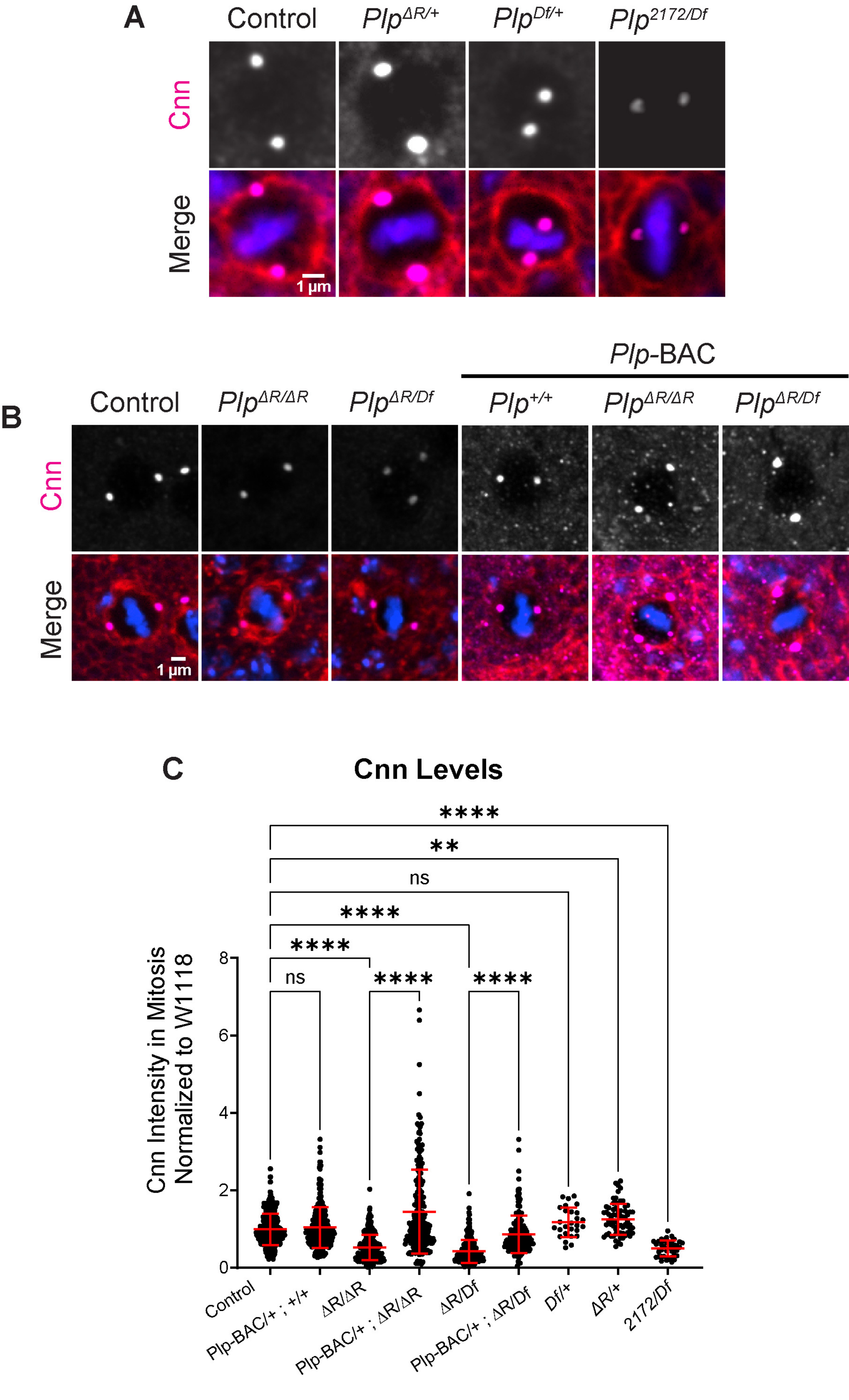


Thomas and Galletta et al., Figure S8

**Figure S8: *plp*$^{\Delta R}$ impacts PCM recruitment.**

Figure is supplement to Fig. 5B, C. Some images and measurements have been duplicated from Fig. 5B, C for ease of comparison. **A-B.** Metaphase wing disc epithelial cells stained for PCM component, Centrosomin (Cnn, magenta)**,** phalloidin (red) and DAPI (blue). **C.** Cnn intensity at metaphase centrosomes normalized to $w^{1118}$ controls. Numbers of samples measured are in Supplementary File 1. n.s.= not significant, ****p ≤ 0.0001.

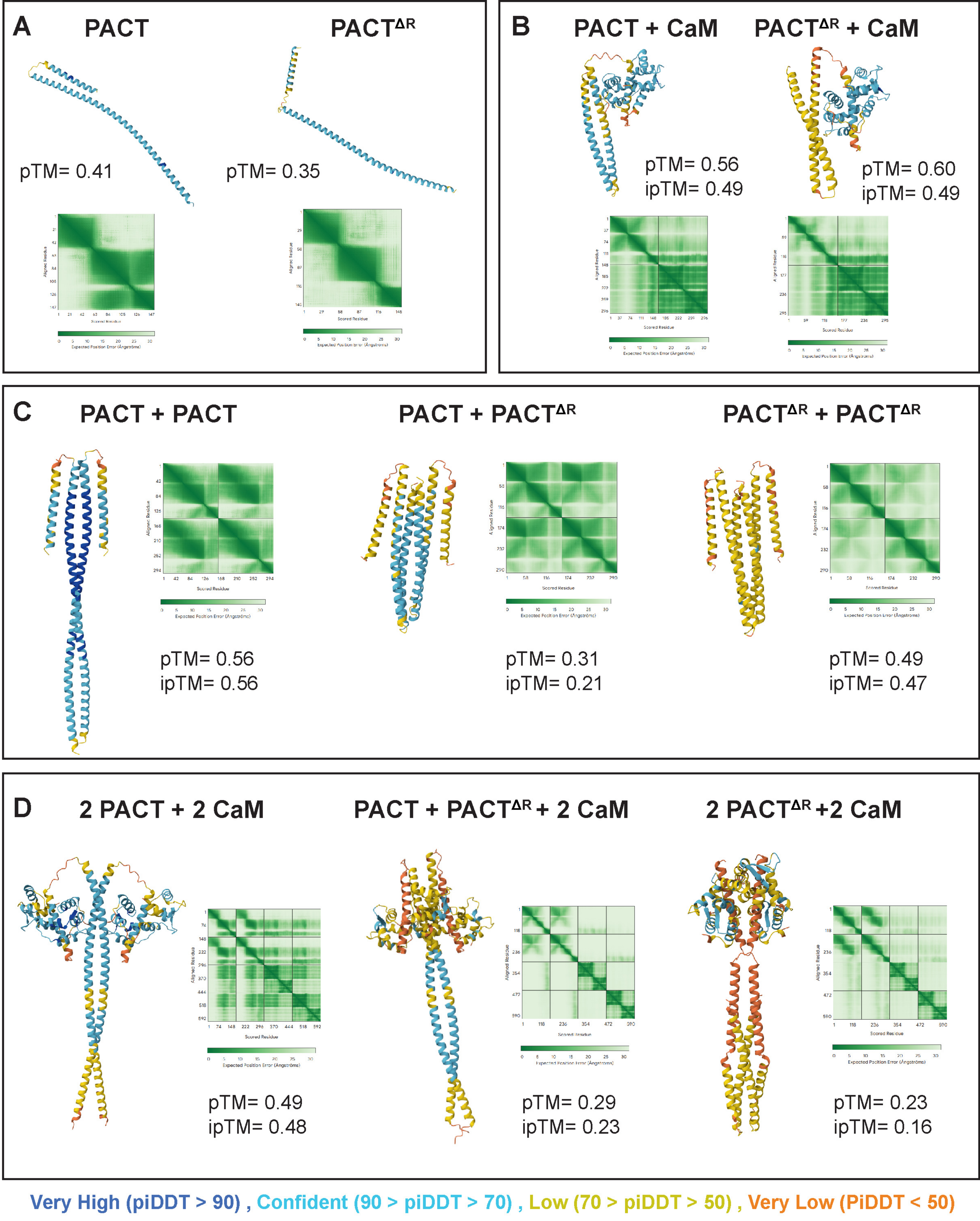


Thomas and Galletta et al., Figure S9

**Figure S9: AlphaFold3 structural predictions of DmPACT.**

Figure is supplement to Figure 6A-C. **A.** Structural predictions of $DmPACT^{WT}$ or $DmPACT^{\Delta R}$ monomers. **B.** Structural predictions of $DmPACT^{WT}$ or $DmPACT^{\Delta R}$ monomers with Calmodulin **C.** Structural predictions of $DmPACT^{WT}$ and $DmPACT^{\Delta R}$ homo and heterodimers. **D.** Structural predictions of 2 $DmPACT^{WT}$ or 2 $DmPACT^{\Delta R}$ with 2 Calmodulin homo and heterodimers. Structures of 2 $DmPACT^{WT}$ or 2 $DmPACT^{\Delta R}$ are reproduced from Figure 7 to aid the reader. pTM and ipTM scores as well as PAE plots included for each structure. Color overlay corresponds to AlphaFold3 confidence scores.

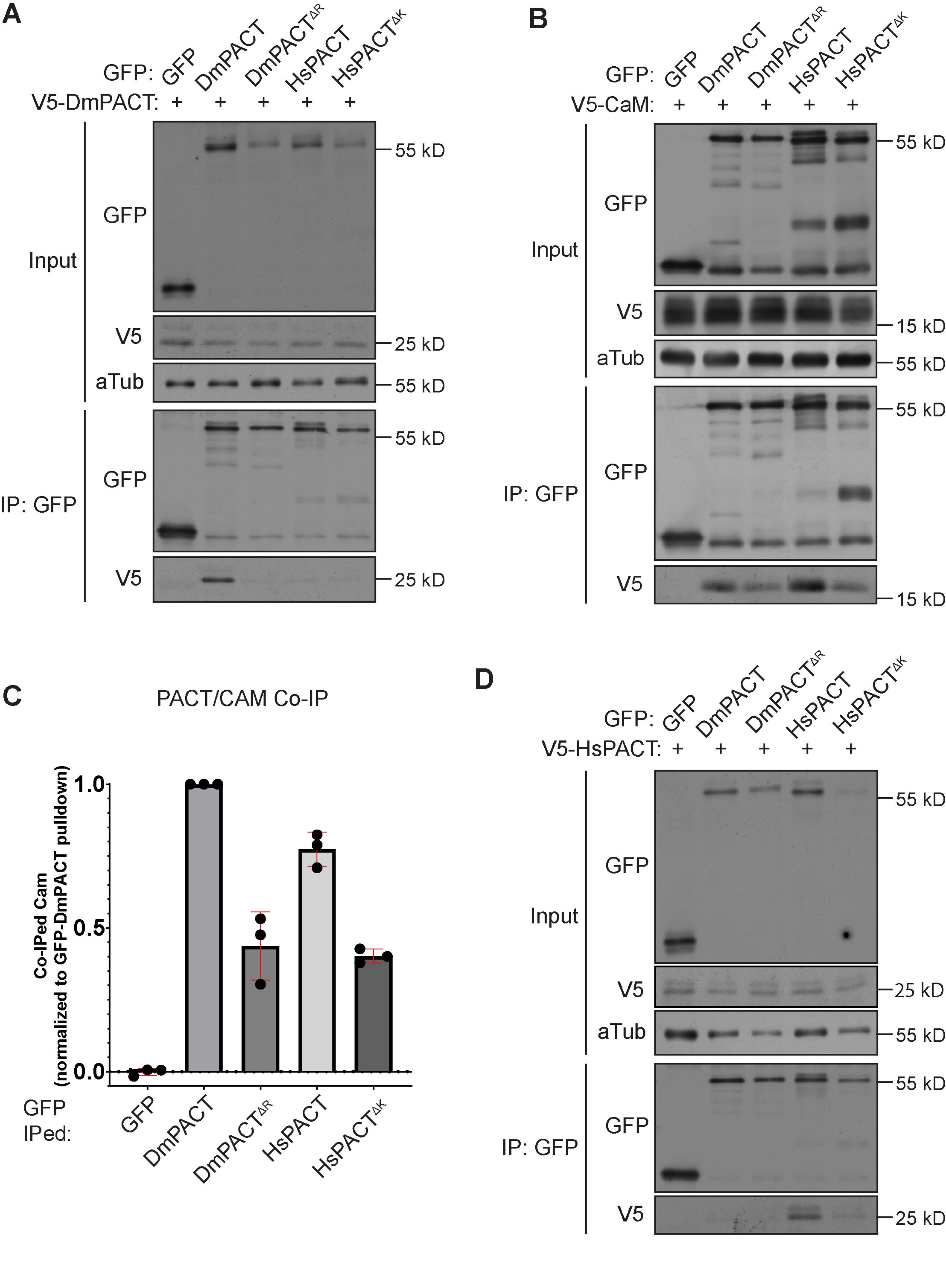


Thomas and Galletta et al., Figure S10

**Figure S10:** ***plp***$^{\Delta R}$ **affects protein-protein interactions.**

Figure is supplement to Figure 6D-F. **A.** Complete co-immunoprecipitation experiment showing DmPACT interactions (Figure 6D) **B.** Complete co-immunoprecipitation experiment showing CaM interactions (Figure 6E). **C.** Quantification of CAM Co-IPed with PACT (Fig. 6E, S10B). Mean ± standard deviation of three replicates. **D.** Complete co-immunoprecipitation experiment showing HsPACT interactions (Figure 6F).

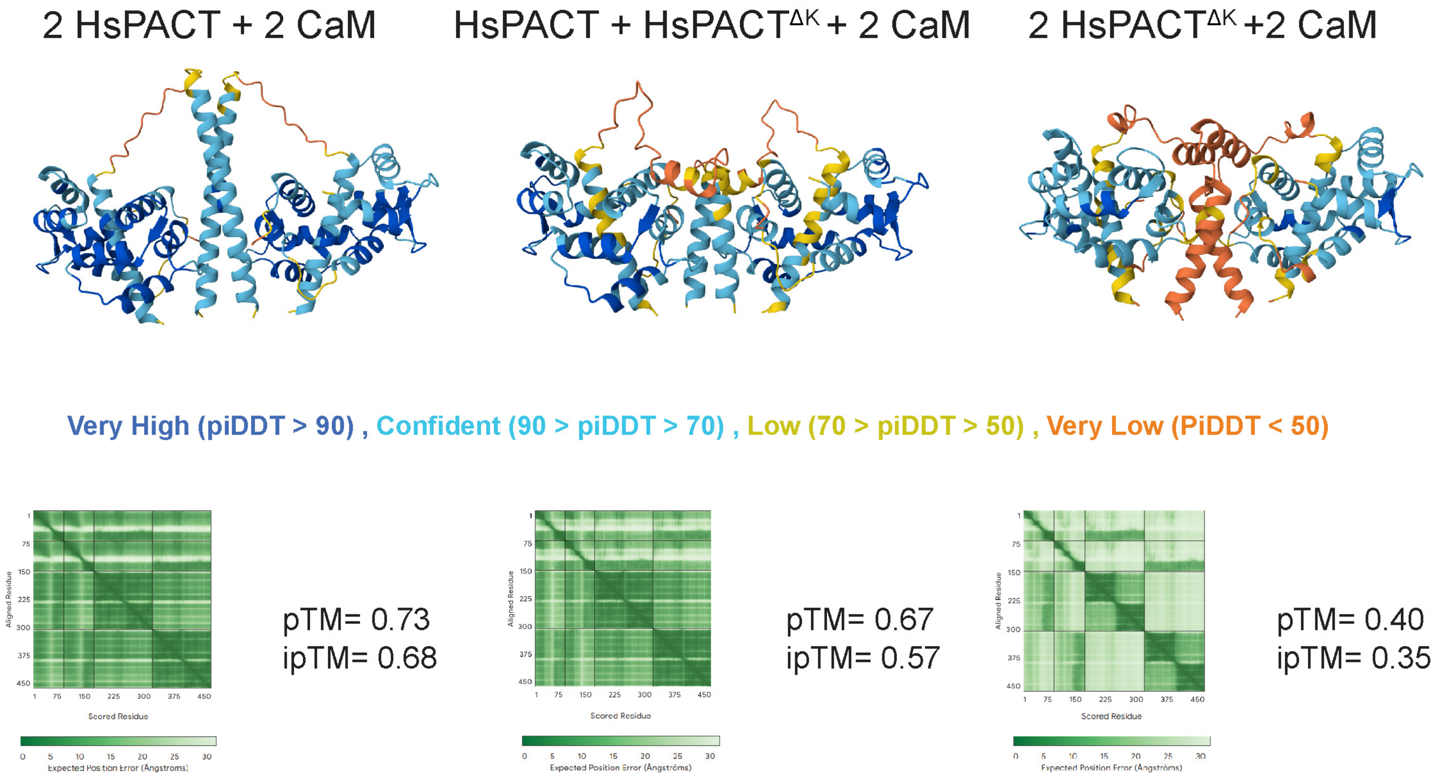


Thomas and Galletta et al., Figure S11

**Figure S11: AlphaFold3 structural predictions of HsPACT.**

Structural predictions of 2 $HsPACT^{WT}$ or 2 $HsPACT^{\Delta K}$ with 2 Calmodulin homo and heterodimers. pTM and ipTM scores as well as PAE plots included for each structure. Color overlay corresponds to AlphaFold3 confidence scores.

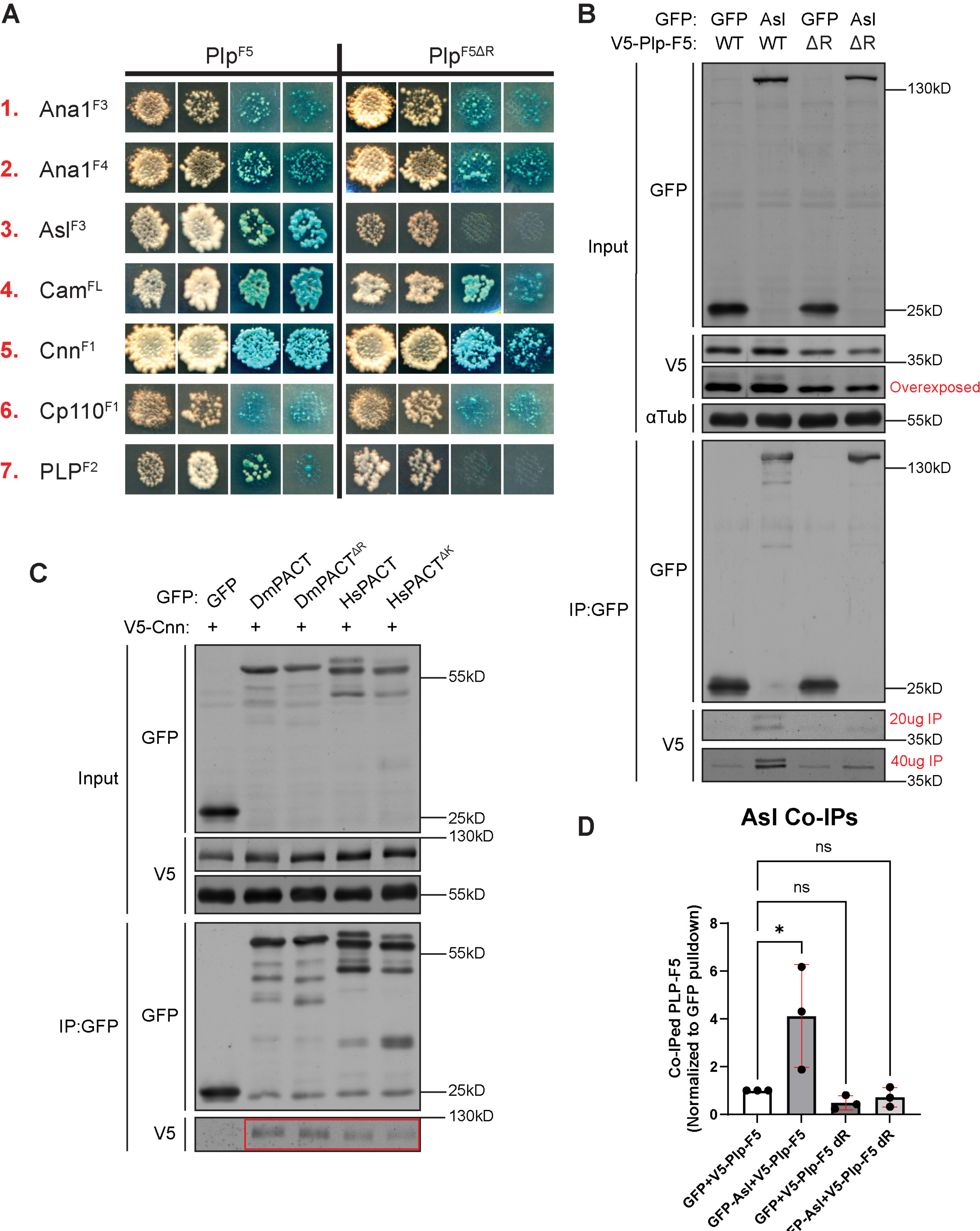


Thomas and Galletta et al., Figure S12

**Figure S12: *plp*$^{\Delta R}$ affects interaction with Asl.**

Figure is supplement to Figure 7. **A.** Complete Y2H experiment depicting interactions of centriole and PCM proteins with PLP(C-term)$^{WT}$ or PLP(C-term)$^{\Delta R}$; the PACT domain resides in this fragment. Images of yeast colonies replica plated on selection plates from left to right: DDO, QDO, QDOX, QDOXA. The DDO and QDOXA iimages are reproduced from Figure 7. **B.** Complete co-immunoprecipitation experiment showing PLP(c-term)/Asterless interactions (Figure 7C). **C**. Co-immunoprecipitation is S2 cells showing V5-Cnn interaction with GFP-DmPACT and GFP-HsPACT, regardless of the presensce of R or K residue. **D.** Quantification of PLP-F5 Co-IPed with Asl (Fig. 7C, S12B). Mean ± standard deviation of three replicates.
